# Engineering of extracellular vesicles for display of protein biotherapeutics

**DOI:** 10.1101/2020.06.14.149823

**Authors:** Dhanu Gupta, Oscar P.B Wiklander, André Görgens, Mariana Conceição, Giulia Corso, Xiuming Liang, Yiqi Seow, Sriram Balsu, Ulrika Felldin, Beklem Bostancioglu, Yi Xin Fiona Lee, Justin Hean, Imre Mäger, Thomas C. Roberts, Manuela Gustafsson, Dara K Mohammad, Helena Sork, Alexandra Bäcklund, C.I. Edvard Smith, Matthew J.A. Wood, Roosmarijn Vandenbroucke, Joel Z. Nordin, Samir EL Andaloussi

**Author notes:** These authors have contributed equally to this work (Dhanu Gupta, Oscar P.B Wiklander, Joel Z. Nordin, Samir EL Andaloussi). Correspondence: Dhanu Gupta, Oscar P.B Wiklander, Joel Z Nordin, Samir EL Andaloussi.

## Abstract

Extracellular vesicles (EVs) have recently emerged as a highly promising cell-free bio-therapeutics. While a range of engineering strategies have been developed to functionalize the EV surface, current approaches fail to address the limitations associated with endogenous surface display, pertaining to the heterogeneous display of commonly used EV-loading moieties among different EV subpopulations. Here we present a novel engineering platform to display multiple protein therapeutics simultaneously on the EV surface. As proof-of-concept, we screened multiple endogenous display strategies for decorating the EV surface with cytokine binding domains derived from tumor necrosis factor receptor 1 (TNFR1) and interleukin 6 signal transducer (IL6ST), which can act as decoys for the pro-inflammatory cytokines TNFα and IL6, respectively. Combining synthetic biology and systematic screening of loading moieties, resulted in a three-component system which increased the display and decoy activity of TNFR1 and IL6ST, respectively. Further, this system allowed for combinatorial functionalization of two different receptors on the same EV surface. These cytokine decoy EVs significantly ameliorated disease phenotypes in three different inflammatory mouse models for systemic inflammation, neuroinflammation, and intestinal inflammation. Importantly, significantly improved *in vitro* and *in vivo* efficacy of these engineered EVs was observed when compared directly to clinically approved biologics targeting the IL6 and TNFα pathways.

## Introduction

Extracellular vesicles (EVs) hold great potential as therapeutic agents with the ability to functionally deliver therapeutic cargos^1^. Our group and others have utilized the display of surface ligands to achieve targeted delivery of nucleic acid species in hard-to-reach tissues, such as the central nervous system (CNS)^2–5^. While being a highly promising strategy, recent studies have highlighted the limitations associated with conventional endogenous surface display technologies, as they typically label only a fraction of the EV population and thus limit the targeting capabilities to a minor sub-set of EVs. Emerging evidence indicates that EVs have numerous subpopulations aside from the classical division into exosomes, microvesicles, and apoptotic bodies^6–9^. This heterogeneity is critically important in EV engineering, especially when delivery of a therapeutic cargo is required in combination with a targeting ligand approach for successful therapy.

Here, we present a novel strategy to display different protein therapeutics simultaneously on the surface of EVs based on synthetic biology and a systematic screening of loading moieties. As proof-of-concept, we targeted inhibition of IL6 and TNFα signaling pathway using an extracellular decoy strategy. Various studies have emphasized that both cytokines play a key role in stimulating inflammation and tissue damage^10,11^. Hence, these pathways are correspondingly targeted by clinically used drugs, including blockers of TNF-receptor (TNFR) (Etanercept, Infliximab) and IL6 receptor (IL6R)(Tocilizumab), to alter the adaptive immune response in autoimmune and inflammatory diseases^12,13^. The soluble TNFα homotrimers exert diverse biological functions, such as cell proliferation, differentiation, and apoptotic signaling, through binding to one of its two receptors, TNFR1 and TNFR2^14^. The cytokine IL6 has broad, pleiotropic biological activities and has been shown to exert both anti-inflammatory and pro-inflammatory signals in deregulated adaptive immune responses^15^. Studies have highlighted that the trans-signaling activation by IL6 complexed to soluble IL6R through IL6 signaling transducer (IL6ST), is linked to inflammation, whereas classical IL6 *cis*-signaling has been shown to be anti-inflammatory and involved in regenerative processes^12^. In this study, we thus aimed to express TNFR and IL6ST on EVs as a clinically relevant approach that enable us to assess the display of the therapeutic proteins on a functional level rather than the mere presence on EV surfaces. Furthermore, the therapeutic relevance of these two cytokines in various inflammatory diseases allowed us to investigate the potency of these receptor decoy systems *in vivo*. Here, a screen of multiple endogenous display strategies was conducted for the decoration of the EV surface with cytokine binding domains of TNFR1 and IL6ST, which can decoy the pro-inflammatory cytokines TNFα and IL6, respectively.

This approach allows us to display more than one receptor type simultaneously in multimeric form and subsequently enhance their inhibitory activity as compared to conventional therapeutics against the same cytokines. In addition, this platform elicits efficient anti-inflammatory effects *in vivo* by significantly improving survival and disease phenotype in three inflammatory mouse models: lipopolysaccharide (LPS)-induced systemic inflammation, experimental autoimmune encephalomyelitis (EAE), and 2,4,6-Trinitrobenzenesulfonic acid (TNBS)-induced colitis, which mimic sepsis, multiple sclerosis (MS), and inflammatory bowel disease (IBD) respectively. The versatility of this engineering approach was further confirmed by successful treatment of EAE using EVs displaying an IL23 alphabody. This work shows great promise for developing engineered, combinatorial EV-based protein therapeutics, as the flexibility of this platform allows robust and efficient surface display of therapeutic proteins and potential targeting ligands.

## Results

### Novel strategies for surface display of biologics on EVs

To develop an efficient EV surface display technology, which can be adapted for targeting domains or therapeutic proteins, we designed numerous surface display designs using luminal EV proteins found to be highly enriched in EV proteomic data sets published by us and others^16– 18^. As a proof-of-concept model for the display of therapeutic proteins on EVs, we fused these EV domains to the cytokine binding domains of either TNFR1 or to IL6ST, for decoy sequestration of TNFα or IL6/IL6R heterodimeric complexes respectively, and further engineered these receptors to be signaling incompetent. This enabled evaluation of various surface display designs in a semi-high throughput workflow by assessing the ability of engineered EVs to decoy their respective cytokines (see schematic illustration of decoy EVs in Figure 1A). An array of genetic constructs were designed using different exosomal sorting proteins, or their respective domains annotated for EV sorting (Figure 1B-E).

**Figure 1.**
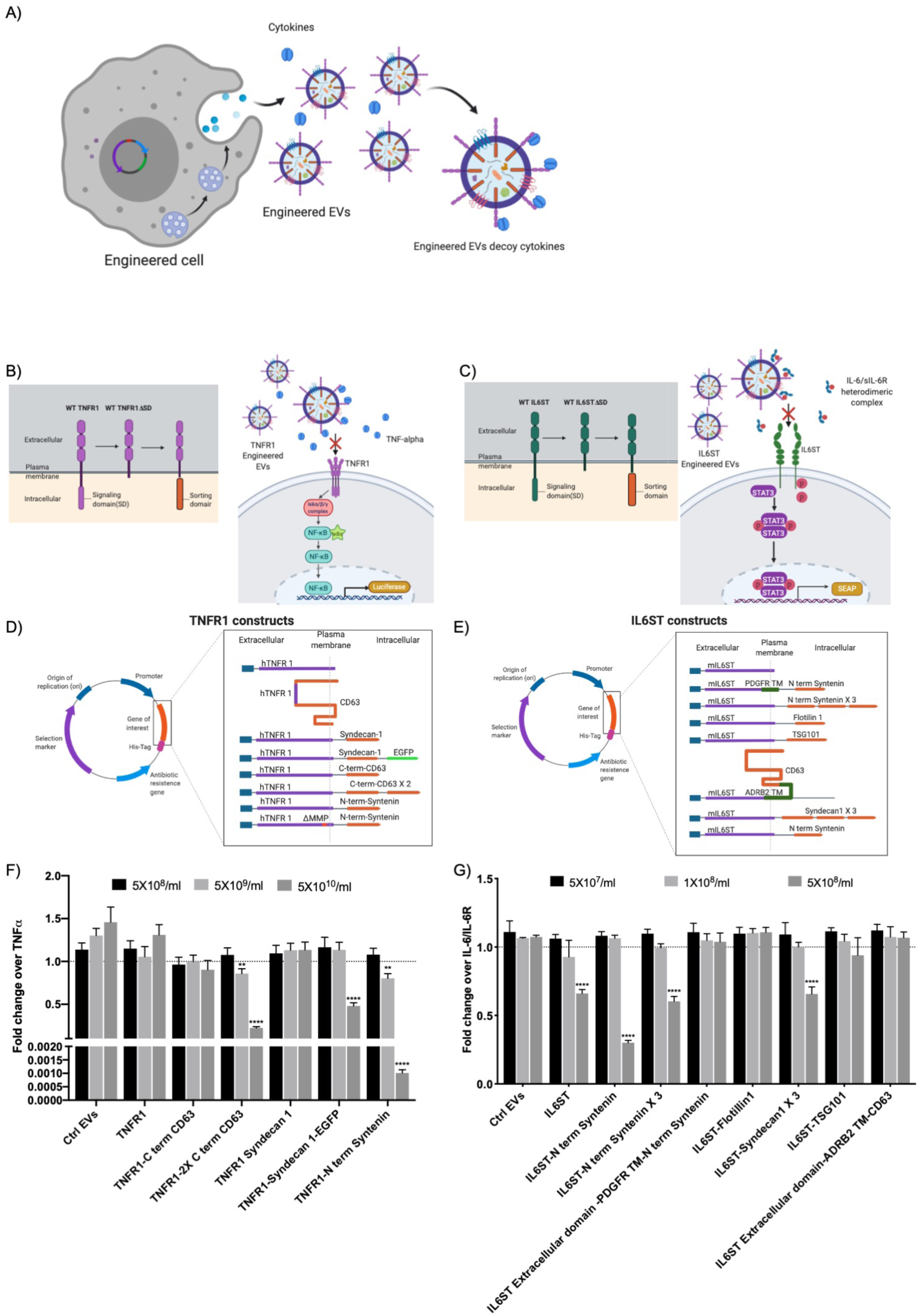
Systematic screening of multiple endogenous EV display strategies for cytokine decoys. **A)** Schematic illustration showing the generation of engineered decoy EVs at the cellular level. Producer cells are genetically modified to express cytokine receptors without the signalling domain fused to an EV sorting domain for efficient display of cytokine receptors on the surface of the secreted EVs (decoy EVs), which can decoy cytokines specifically. **B) and C)** Schematic illustrations of the evolution of cytokine receptors to facilitate EV surface display and assessment of various designs in a cytokine induced reporter cell system for high throughput screening of TNFR1 and IL6ST decoy EVs. **D) and E)** List of various TNFR1 or IL6ST sorting domain fusions assessed in the initial screen. **F)** Engineered decoy EVs displaying TNFR1 purified from HEK293T cells transfected with the constructs encoding the different display constructs (Figure1D) evaluated for TNFα decoy in an *in vitro* cell assay responsive to TNFα induced NF-κB activation. Data were normalized to control cells treated with TNFα only (5 ng/ml). **G)** Engineered EVs displaying IL6ST purified from HEK293T cells transfected with constructs encoding the different display constructs (Figure1G) evaluated for IL6/sIL6R decoy in an *in vitro* cell assay respondent to IL6/sIL6R induced STAT3 activation. Data were normalized to control cells treated with IL6/sIL6R (5 ng/ml). **F, G**, Error bars, s.d. (*n* = 3), **** *P* < 0.0001, *** *P* < 0.001, ** *P* < 0.01, statistical significance calculated by two-way ANOVA with Dunnett’s post-test compared with response of Ctrl EVs at the respective dose.

HEK293T cells were transiently transfected with plasmids encoding the different display constructs and engineered EVs were purified by ultracentrifugation and quantified by nanoparticle tracking analysis (NTA) (Supp. Figure 1A and 1B). The potency of the purified EVs was assessed using an *in vitro* reporter system for the respective cytokine, either by detecting luciferase activity driven by a NF-κB minimal promoter (TNFα, Figure 1B and Supp. Figure 1C) or secreted alkaline phosphatase (SEAP) driven by a STAT3 minimal promoter (IL6/sIL6R, Figure 1C and Supp. Figure 1D). We observed inhibitory activity of engineered decoy EVs with various designs in a dose-dependent manner (Figure 1F, G). Constructs with inclusion of the N-terminal sorting domain derived from Syntenin (TNFR1-N term Syntenin and IL6ST-N term Syntenin), a protein implicated in sorting of protein cargo into EVs, significantly and reproducibly exhibited the best inhibitory activity for both IL6ST and TNFR1 signaling incompetent receptor constructs (Figure 1D-G and Supp. Figure 2A). Furthermore, the functionality of the cytokine decoy EVs was corroborated by quantitative assessment of EVs by western blot (WB) probing for the respective decoy receptor (TNFR1 or IL6ST) or the fused His-Tag on C terminus of each construct. Notably, TNFR1-N term Syntenin (Supp. Figure 2B & 3) and IL6ST-N term Syntenin (Supp. Figure 4 & 5) displayed a clear band in both parental cells and EVs of respective decoy receptor. Interestingly, we observed cleavage products of the TNFR1 fusion protein only upon probing the WB with the His-Tag antibody and not with the hTNFR1 antibody. We hypothesized that a matrix metalloprotease (MMP) cleavage site previously annotated^19^ in TNFR1 resulted in cleavage of the receptor from the decoy EV surface, hence reducing the efficacy of the TNFR1-decoy EVs. Deletion of the MMP cleavage site on the TNFR1 extracellular domain reduced the levels of the cleaved product and resulted in a further 10-fold increase of the inhibitory activity in reporter cells (Supp. Figure 6A-B).

To further increase efficiency, multimerization domains were introduced in different positions within the constructs to increase the number of decoy receptors per EV and to mimic the natural receptor state *in situ*^12,13^. A trimerization domain ‘Foldon’ derived from T4 fibritin protein of T4 bacteriophage^20^ was introduced to the lead TNFR1 design either in the extracellular- or intracellular region (Figure 2A). The addition of a multimerization domain to the TNFR1-N term Syntenin construct further increased the efficiency of the decoy EVs to sequester TNFα (Figure 2B-C). Similarly, we introduced a dimerization domain ‘GCN4 L.Z’, derived from yeast^21^, and a tetramerization domain ‘Fragment X’, derived from Phosphoprotein P of human metapneumovirus^22^, to the IL6ST-N term Syntenin construct in the intracellular domain (Figure 2D). Both designs showed a significant enhancement over their predecessors (Figure 2E, F). However, addition of the dimerization domain both in the extracellular and intracellular domain decreased the efficacy of the display, potentially because it affects the cytokine binding properties of the receptor. Furthermore, a sorting domain derived from transferrin receptor (TfR) along with GCN4 L.Z dimerization domain showed similar efficacy as compared to Syntenin in our screen when it was fused to IL6ST. Importantly, the variation in active dose among the different sets of experiments is primarily due to large-scale transfection and therefore a relative comparison with multiple doses was performed in every screen to account for this variation.

**Figure 2.**
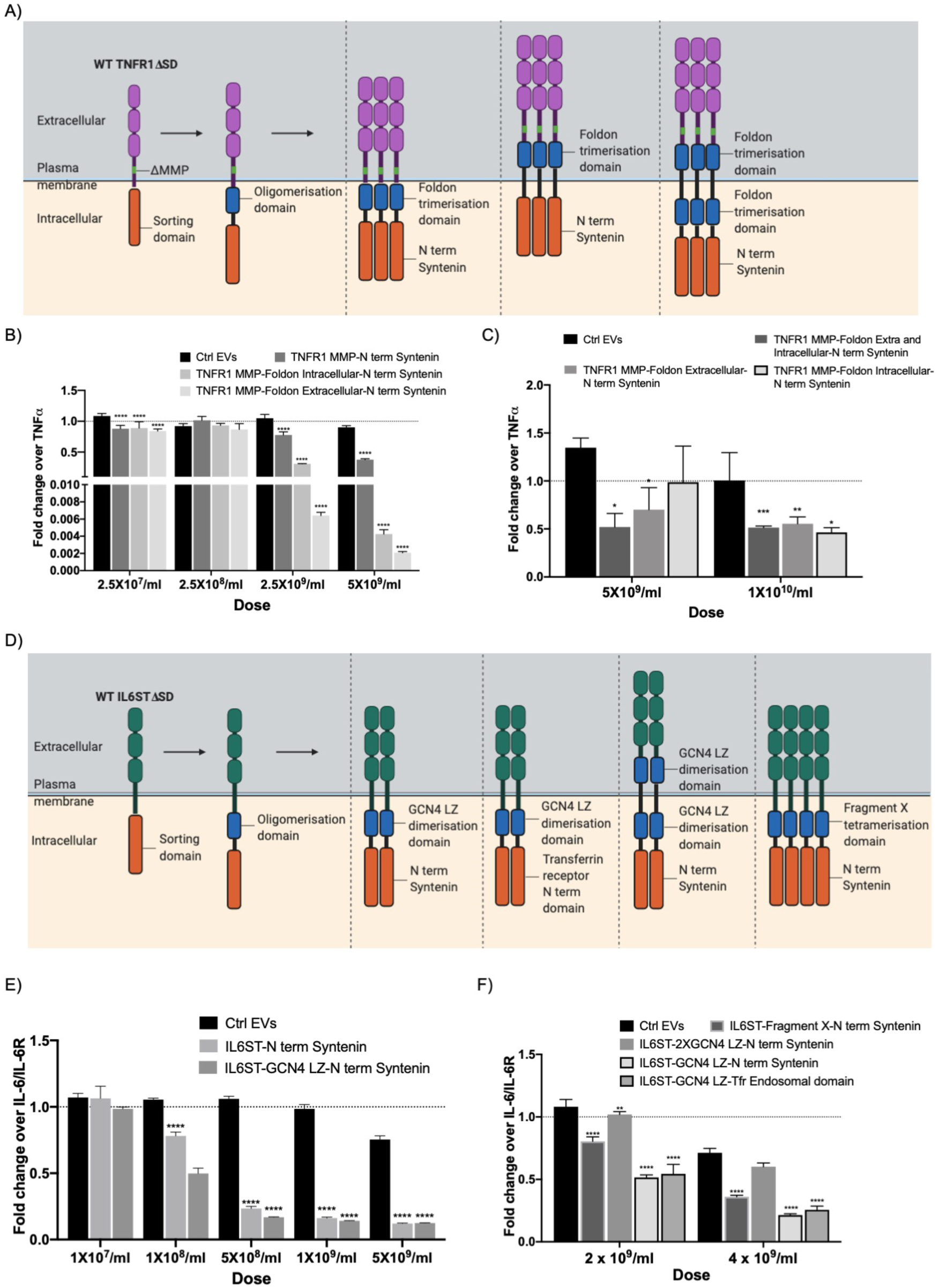
Multimerization of decoy fusion proteins improves loading and activity. **A)** Schematic illustration showing the evolution of TNFR1 design by addition of trimerization domains to enhance loading and binding efficiency of the EV- displayed decoy receptors. **B) and C)** Systematic comparison of various TNFR1 designs with multimerization domains. Engineered EVs displaying TNFR1 purified from HEK293T cells transfected with constructs encoding TNFR1 multimerization sorting domain fusion proteins (as listed in Figure 2A) evaluated for TNFα decoy in an *in vitro* cell assay responsive to TNFα induced NF-κB activation. Data were normalized to control cells treated with TNFα only (5 ng/ml). **D)** Schematic illustration showing the evolution of IL6ST designs by addition of different multimerization domain to enhance loading and binding efficiency of displayed decoy receptors on EVs. **E) and F)** Engineered EVs displaying IL6ST purified from HEK293T cells transfected with constructs encoding IL6ST multimerization sorting domain fusion constructs (as listed in Figure 2D) respectively evaluated for IL6/sIL6R decoy in an *in vitro* cell assay respondent to IL6/sIL6R induced STAT3 activation. Data were normalized to control cells treated with IL6/sIL6R (5 ng/ml). **B, C, E, F**, Error bars, s.d. (*n* = 3), **** *P* < 0.0001, *** *P* < 0.001, ** *P* < 0.01, * *P* < 0.05, statistical significance calculated by two-way ANOVA with Dunnett’s post-test compared with response of Ctrl EVs at the respective dose.

Based on these findings, we selected N-term-Syntenin as the EV-sorting domain for both TNFR1 and IL6ST decoy EVs with the intracellular dimerization domain included for IL6ST (IL6STΔ-LZ-NST) and with the intracellular trimerization domain and deletion of MMP-cleavage site for TNFR1 (TNFR1ΔΔ-FDN-NST) (Figure 2A and 2D). As foreign protein sequences in the extracellular domains might lead to immune responses that could affect multi-dosing strategies, designs with intracellular multimerization domains were chosen for subsequent work^23^.

To reduce the variability associated with cellular transfection and to further scale up the production of therapeutic EVs for *in vivo* applications, stable engineered HEK293T producer cells for production of IL6STΔ-LZ-NST- and TNFR1ΔΔ-FDN-NST-decoy EVs were established using lentiviral transduction. HEK293T IL6STΔ-LZ-NST decoy EVs inhibited IL6/IL6R induced trans-signaling with reduced STAT3 activation and TNFR1ΔΔ-FDN-NSTz-decoy EVs could inhibit TNFα stimulated NF-kB activation in a dose dependent manner (Supp. Figure 7A and 7B). To further validate whether our EV engineering strategy is applicable to other cell types, we validated the approach in a more therapeutically relevant cell source, mesenchymal stem cells (MSC), which were engineered to stably produce the respective decoy EVs (Supp. Figure 7C). MSC TNFR1ΔΔ-FDN-NST and IL6STΔ-LZ-NST EVs displayed a dose response to decoy the cytokines similar to what was observed using HEK293T cell-derived decoy EVs (Figure 3A-B). The engineered MSC-derived EVs were characterized using NTA, showing that the majority of the EVs were in the size range of exosomes with a peak of around 100 nm (Supp. Figure 7D). Characterization by WB of isolated EVs from the respective cell source confirmed expression of both common EV markers ALIX and TSG101, absence of apoptotic body marker calnexin and presence of the respective decoy proteins (Figure 3C and Supp. Fig 8)^24^. In addition, the presence of decoy receptors (TNFR1 or IL6ST) on EVs was confirmed by immunogold electron microscopy, using primary antibodies against the respective decoy receptor (Figure 3D).

**Figure 3.**
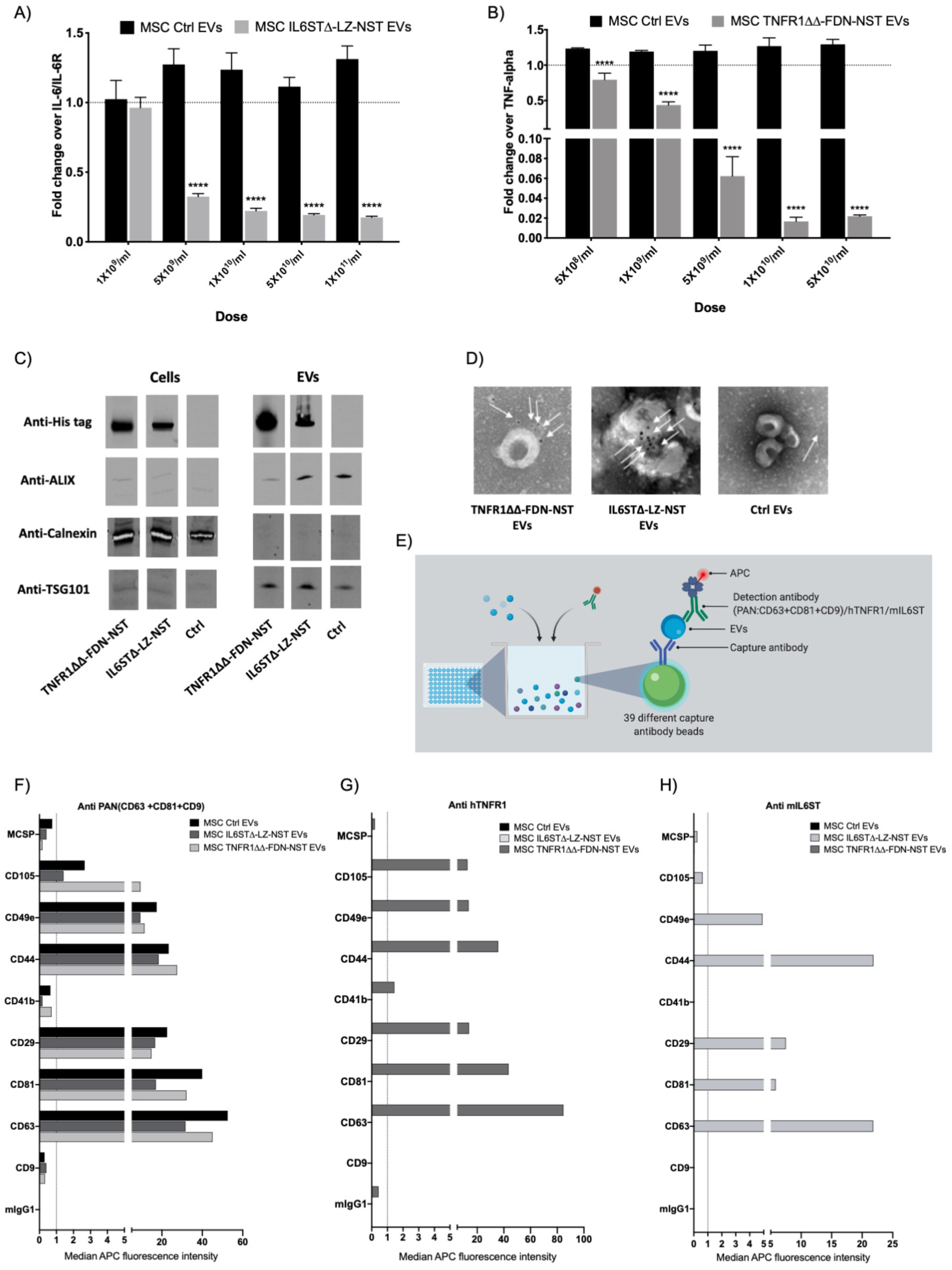
Multimeric decoy receptor EV sorting protein chimera is functionalised on several EV subpopulations. **A)** Engineered EVs displaying IL6ST purified from MSC cells stably expressing the optimised IL6STΔ-LZ-NST display construct, evaluated for IL6/sIL6R decoy in an *in vitro* cell assay respondent to IL6/sIL6R induced STAT3 activation. EVs purified from MSC stably expressing Ctrl construct were used as control. Data were normalized to control cells treated with IL6/sIL6R (5 ng/ml). **B)** Engineered EVs displaying TNFR1 purified from MSC cells stably expressing the optimized TNFR1ΔΔ-FDN-NST display construct, evaluated for TNFα decoy in an *in vitro* cell assay responsive to TNFα induced NF-κB activation. EVs purified from MSC stably expressing Ctrl construct were used as control. Data were normalized to control cells treated with TNFα (5 ng/ml) treated cells. **C)** WB of MSC TNFR1ΔΔ-FDN-NST, IL6STΔ-LZ-NST and Ctrl cells and EVs indicating the presence of classical EV markers; ALIX (96 kDa), TSG101 (44 kDa) and absence of Calnexin (67 kDa) in the isolated EVs. The WB results further demonstrate the presence of respective His-tagged decoy receptors; TNFR1ΔΔ-FDN-NST (48 kDa) and IL6STΔ-LZ-NST (94 kDa) both on cells and EVs. **D)** Transmission electron microscopy of MSC TNFR1ΔΔ-FDN-NST, IL6STΔ-LZ-NST and Ctrl-EVs with nanogold labelled antibody staining of respective decoy receptor indicated by white arrows. **E)** Schematic illustration showing the workflow of the multiplex bead-based flow cytometry assay. Isolated EVs incubated with up to 39 different bead populations coated with different capture antibodies, which are distinguishable by flow cytometry due to their different fluorescence intensities. EVs captured by the different beads are detected with detection antibodies either against PAN (CD63-APC, CD81-APC and CD9-APC), mIL6ST-APC, or hTNFR1-APC. **F-H)** Characterization of EV surface protein composition by using F) anti-PAN (CD63, CD81 and CD9), G) anti-hTNFR1 and H) anti-mIL6ST detection antibodies in multiplex bead-based assays to confirm marker co-expression on MSC TNFR1ΔΔ-FDN-NST, IL6STΔ-LZ-NST and ctrl EVs. Data represented as background corrected median APC fluorescence intensity determined by flow cytometry of EVs bound to respective capture beads and upon using APC labelled detection antibody. **A, B**, Error bars, s.d. (*n* = 3), **** *P* < 0.0001, statistical significance calculated by two-way ANOVA with Dunnett’s post-test compared with response of Ctrl EVs at the respective dose.

To determine the impact of this engineering strategy on the EV surface proteomic profile, a multiplex bead-based flow cytometry assay was applied for simultaneous flowcytometric detection of 37 surface proteins on CD63/CD81/CD9 positive vesicles^25^ (Figure 3E). MSC-derived TNFR1ΔΔ-FDN-NST and IL6STΔ-LZ-NST decoy EVs exhibited a highly similar surface protein profile as compared to MSC Ctrl EVs for 37 different surface markers on tetraspanin positive vesicles (Figure 3F and Supp. Figure 9A-C). Furthermore, to determine whether the decoy receptors are present on the tetraspanin positive subpopulation of EVs, we modified the bead-based assay by using decoy receptor-based detection instead of tetraspanin-based detection of the 37 different capture beads. Upon using hTNFR1 antibody as a detection antibody for assessing the 37 different antigens, CD63 and CD81 were observed to be the most enriched surface markers on TNFR1ΔΔ-FDN-NST positive vesicles, whereas Ctrl EVs and IL6STΔ-LZ-NST EVs were negative for all markers (Figure 3G and Supp. Figure 9D-F). A similar trend was observed for IL6STΔ-LZ-NST EVs upon using mIL6ST antibody as a detection antibody for the capture beads (Figure 3H and Supp. Figure 9G-I). These results clearly indicate that our engineering strategy allows for highly efficient engineering of EVs with negligible disruption of their endogenous surface protein profile.

### Engineered decoy EVs display improved efficacy compared to conventional biologics

Next, we sought to investigate the efficacy of engineered EVs and compare them to a clinically approved biologic against TNFα, Etanercept (a dimeric TNFR2 protein). Using the aforementioned TNFα reporter model, HEK293T TNFR1ΔΔ-FDN-NST decoy EVs showed a 10-fold lower IC50 value compared to Etanercept^13^ (Figure 4A-B). Moreover, in order to further validate the potency of engineered EVs, a physiologically more relevant *in vitro* model of inflammation was utilized. Using RAW 246.7 macrophages challenged with LPS, HEK293T TNFR1ΔΔ-FDN-NST decoy EVs significantly downregulated secreted TNFα levels in RAW 246.7 macrophages, in a dose-dependent manner both at 6 hours and 24 hours post stimulation (Figure 4C and Supp. Figure 10). After these encouraging *in vitro* results, the potency of decoy EVs was assessed *in vivo* using an LPS-induced systemic inflammation mouse model. Intravenous (IV) injection of HEK293T decoy EVs along with LPS challenge in mice resulted in significantly improved survival (100% up to 60 hours) of TNFR1ΔΔ-FDN-NST- and IL6STΔ-LZ-NST EV treated mice compared to 0% survival of mock treated mice (Figure 4B). This was further corroborated with MSC-derived TNFR1ΔΔ-FDN-NST decoy EVs (1×10^11^), which showed improved survival (100% up to 60 hours) compared to 160 *µ*g Etanercept (25% at 60 hours) and 1×10^11^ Ctrl EVs (50% at 60 hours) (Figure 4E). In a separate experiment, mice treated with 6.5×10^11^ MSC IL6STΔ-LZ-NST and/or 6.5×10^11^ TNFR1ΔΔ-FDN-NST decoy EVs displayed reduced weight loss (Figure 4F). The protective effect in mice with LPS-induced inflammation was further improved by combinatorial treatment with both decoy EVs, as compared to 6.5×10^11^ unmodified MSC-EVs. These results collectively underpin the therapeutic potential of decoy EVs and led us to continue assessing the therapeutic anti-inflammatory potential of decoy EVs in an autoimmune MS disease model.

**Figure 4.**
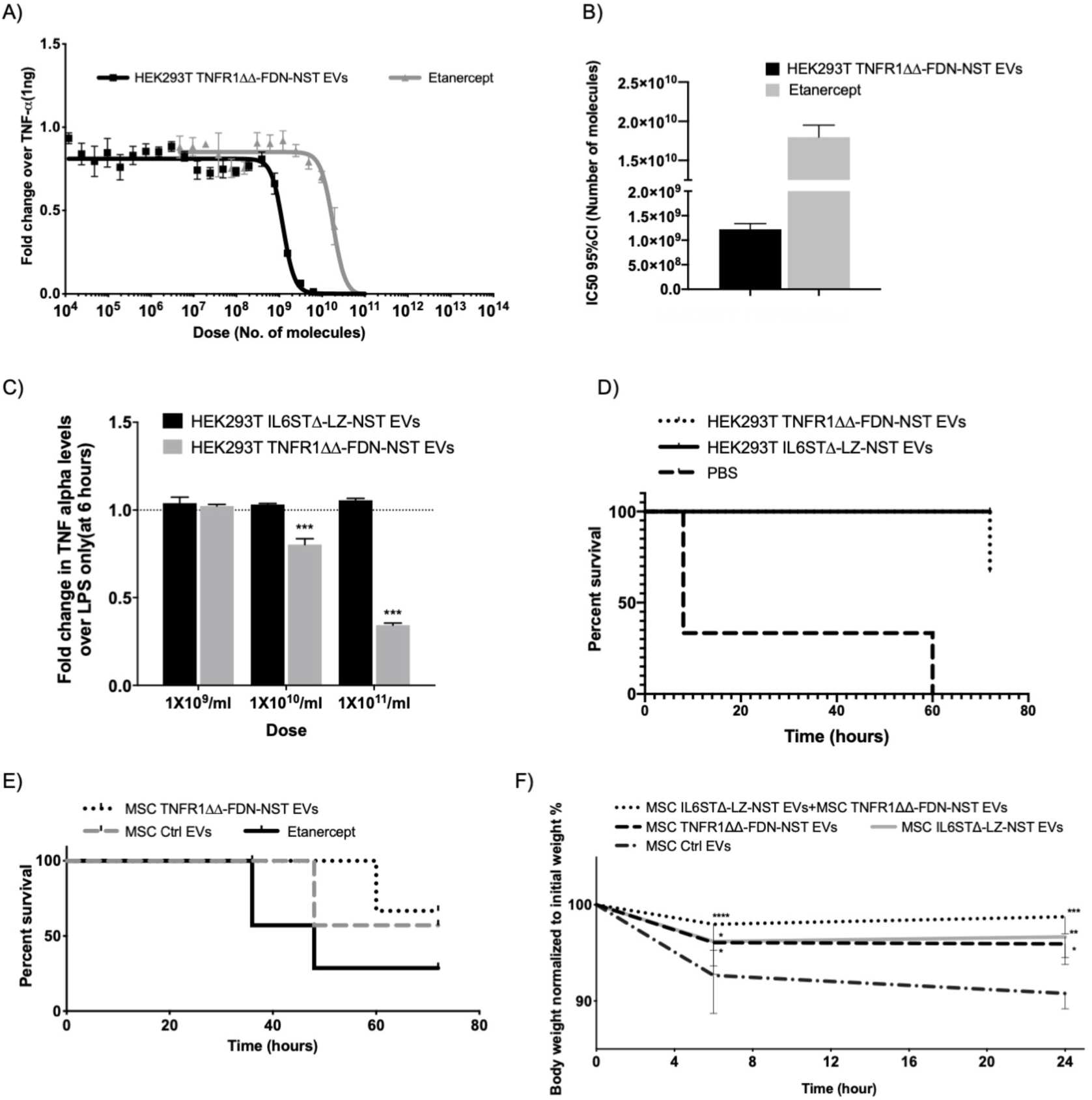
Benchmarking Engineered decoy EVs against clinically approved biologics both *in vitro and in vivo*. Comparison of **A)** inhibitory dose response curves and **B)** calculated IC50 values (95% Confidence interval) for TNFα sequestration by HEK293T TNFR1ΔΔ-FDN-NST EVs and Etanercept. HEK293T NF-kB reporter cells were challenged with 10 ng/ml (1 ng) of TNFα along with increasing doses of either HEK 293T TNFR1ΔΔ-FDN-NST EVs or Etanercept. **C)** Effect of HEK293T TNFR1ΔΔ-FDN-NST EVs and IL6STΔ-LZ-NST EVs on TNFα levels in conditioned medium determined by ELISA at 6 hours post LPS stimulation of RAW 246.7 macrophages. Data were normalized to control cells treated with LPS only. **D)** Survival curve of LPS (15 mg/kg) induced systemic inflammation in mice treated with intravenous injection of either 1×10^11^ HEK 293T TNFR1ΔΔ-FDN-NST EVs (*n*=3) or 2×10^11^ HEK293T IL6STΔ-LZ-NST EVs (*n*=4) or PBS (*n*=5) 3 hours post induction. **E)** Survival curve of LPS (15 mg/kg) induced systemic inflammation in mice treated with intravenous injection of either 1×10^11^ MSC TNFR1ΔΔ-FDN-NST EVs or 1×10^11^ MSC Ctrl EVs or 160 *µ*g Etanercept 3 hours post LPS induction. **F)** Percent relative bodyweight to initial bodyweight over time of mice induced with LPS (15 mg/kg). Mice were treated with intravenous injection of either 3.25×10^11^ MSC IL6STΔ-LZ-NST EVs + 3.25×10^11^ MSC TNFR1ΔΔ-FDN-NST EVs (*n*=6), 6.5×10^11^ MSC TNFR1ΔΔ-FDN-NST EVs (*n*=6), 6.5×10^11^ MSC IL6STΔ-LZ-NST EVs (*n*=6), 6.5×10^11^ MSC Ctrl EVs (*n*=6), or PBS (*n*=6) 3 hours post induction. **A**, Error bars, S.E.M (*n*=3), **** *P* < 0.0001, statistical significance calculated by two-way ANOVA with Dunnett’s post-test compared with response of Ctrl EVs at the respective dose. **F**, Error bars, S.E.M (*n* = 6), **** *P* < 0.0001, *** *P* < 0.001, ** *P* < 0.01, * *P* < 0.05, statistical significance calculated by two-way ANOVA with Dunnett’s post-test compared with response of Ctrl EVs at the respective observation time.

### Engineered decoy EVs inhibit progression of neuroinflammation

We have previously shown that EVs can be used to treat hard-to-reach tissues, including the ability to cross the blood-brain-barrier and exhibit therapeutic effects in the Central nervous system(CNS)^4^. To explore the potential effect of decoy EVs in neuroinflammation, the experimental autoimmune EAE mouse model, mimicking MS in humans, was used. To evaluate the effect of treatment in these mice, clinical scores on a scale of 0-5 are used that reflect the disease severity (EAE-score, Table 1). Upon repeat subcutaneous (SC) administration of control MSC EVs, as depicted in Figure 5A, no effect was observed on disease progression compared to mock treatment. However, as hypothesized, TNFR1ΔΔ-FDN-NST decoy EVs (4×10^10^) Subcutaneous (SC) treatment significantly reduced disease progression over time (Figure 5B). At the endpoint (day 16), mice treated with decoy EVs could still move freely (Supp. Movie 1), with only minor tail and/or hind limb weakness, as compared to mock treated mice that were hind limb paralyzed. Additionally, significantly lower EAE-scores were observed in mice treated with TNFR1ΔΔ-FDN-NST decoy EVs (EAE-score 1.7/5) compared to control EVs (EAE score 3.0), which was similar to mock treatment (EAE score 2.9/5) (Figure 5C). In addition, mock treated EAE mice displayed a gradual decrease in body weight after symptom onset, reflecting disease progression (Supp. Figure 11A). Mice treated with TNFR1ΔΔ-FDN-NST decoy EVs displayed sustained bodyweight over time, with increased weight at the endpoint (4.2%) compared to mock treated mice (Supp. Figure 11B). In addition, treatment with TNFR1ΔΔ-FDN-NST decoy EVs reduced levels of the pro-inflammatory cytokines TNFα and IL6 in spinal cord compared to mock treatment (Supp. Figure 12A and 12B).

**Figure 5.**
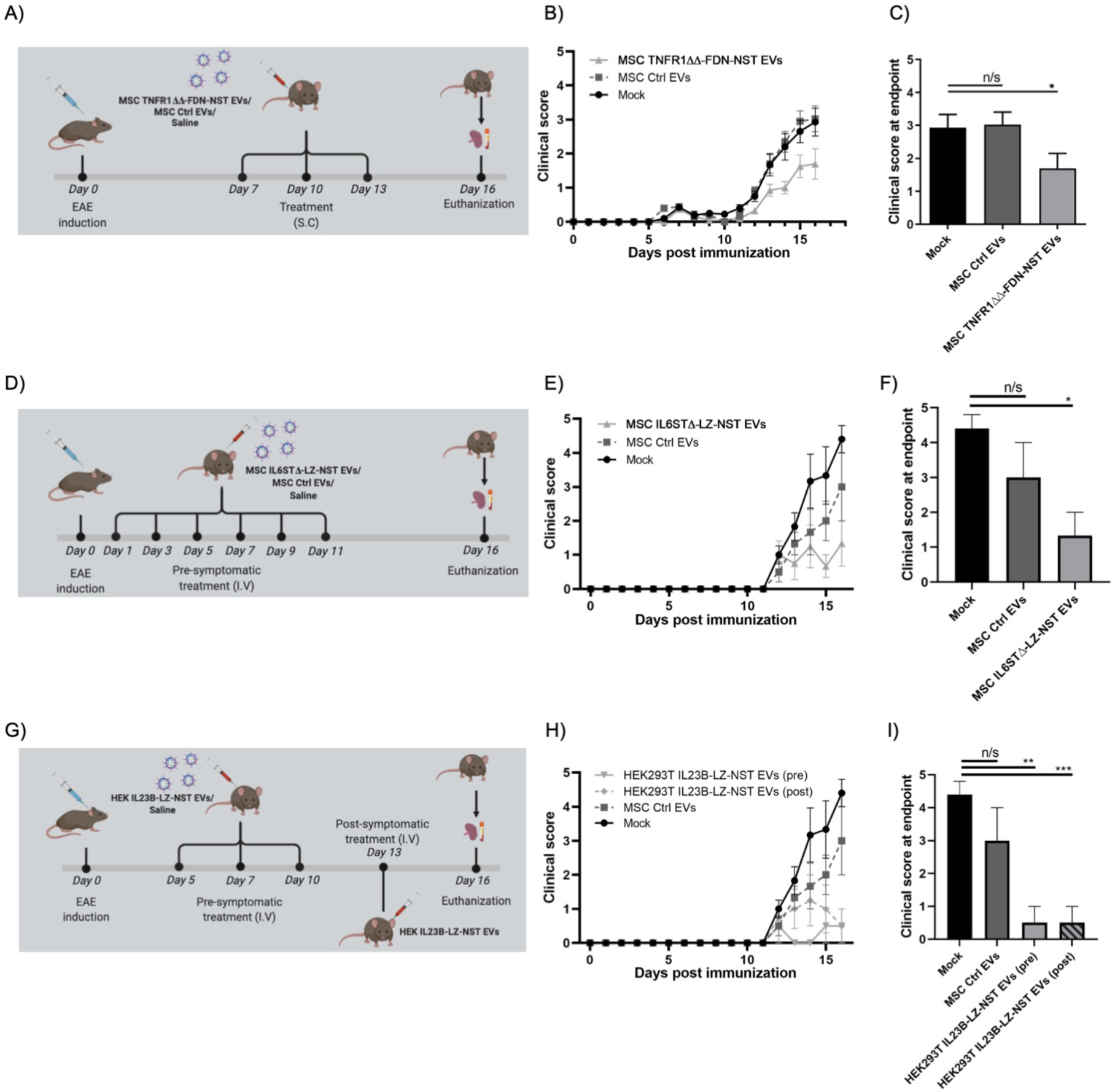
Targeting TNF-alpha, IL6 and IL23 signalling axis with engineered decoy EVs suppress neuroinflammation. A) Description of the treatment protocol for TNFα decoy EVs in EAE. **B)** Clinical score (EAE-score, see Supplementary table 1) of disease progression over time and **C)** EAE-score at endpoint (day 16) in mice induced with EAE using MOG_35-55_ peptide and treated with subcutaneous (S.C) administration of either 4×10^10^ MSC TNFR1ΔΔ-FDN-NST EVs (*n*=5), MSC Ctrl EVs (*n*=5), or saline (*n*=5) (on day 7, 10 & 13). **D)** Schematic description of treatment protocol for IL6 decoy EVs in EAE. **E)** Clinical score of disease progression over time and **F**) EAE-score at endpoint (day 16) in mice induced with EAE using MOG_35-55_ peptide and treated with intravenous (I.V) administration of either 5×10^9^ MSC IL6STΔ-LZ-NST EVs (*n*=5), MSC Ctrl EVs (*n*=5), or saline (*n*=6) (on day 1,3,5,7,8,9 & 11). **G)** Schematic description of treatment protocol for IL23 decoy EVs in EAE. **H)** Clinical score of disease progression over time and **I)** EAE-score at endpoint (day 16) in mice induced with EAE using MOG_35-55_ peptide and treated I.V with either 1×10^10^ HEK293TIL23B-LZ-NST EVs pre symptomatic (*n*=5) (on day 5, 7 & 10), 6×10^10^ HEK293TIL23B-LZ-NST EVs post symptomatic (*n*=5) (on day 13), or saline (*n*=6). **C, F, I**, Error bars, SEM *** *P* < 0.001, ** *P* < 0.01, * *P* < 0.05 statistical significance calculated by two-way ANOVA with Dunnett’s post-test compared with response to mock treated animal.

In a similar set-up, depicted in Figure 5D, we next tested the therapeutic potential of blocking IL6 signaling in neuroinflammation, but instead of a sustained release route we opted for a rapid release route (IV, intravenous) for an immediate effect. Repeated injection of IL6STΔ-LZ-NST EVs (5×10^9^) in mice induced with EAE until onset of symptoms, showed significant reduction in clinical score at day 16 (EAE score 1.33/5) as compared to mock treatment (EAE score 4.4/5) (Figure 5E-F). In contrast, MSC control EVs showed only minimal therapeutic effect (EAE score 3/5). Both TNFR1ΔΔ-FDN-NST and IL6STΔ-LZ-NST decoy EVs thus significantly reduced disease progression in a neuroinflammatory MS model.

To further test the versatility of this display system, we designed constructs using IL6ST-LZ-NST as a backbone and replaced the cytokine binding region of IL6ST with an alphabody against IL23^26^, another pro-inflammatory cytokine implicated in the pathophysiology of MS^27^. Repeated injection of IL23B-LZ-NST EVs (1×10^10^) purified from transiently transfected HEK293T cells, after the induction of disease, showed significantly lower EAE scores (EAE score 0.4/5) (Figure 5G-I and Supp. Figure 12C). Importantly, administration of a single dose of IL23 decoy EVs (6×10^10^) in EAE mice after onset of symptoms reduced the clinical score compared to mock treatment (Figure 5I). Taken together, these data clearly reflect the adaptability of the engineering platform and the potential of using EVs to display therapeutic receptors in inflammatory diseases including hard-to-treat CNS inflammation.

### Double decoy EVs display two receptors simultaneously and effectively abrogate colitis in mice

After successful application of engineered EVs displaying single biologics, we next sought to generate combinatorially engineered EVs displaying two different surface proteins simultaneously. To this end, MSC cells stably expressing both TNFR1ΔΔ-FDN-NST and IL6STΔ-LZ-NST were generated (Supp. Figure 13A). The optimized engineered EVs isolated from conditioned medium were characterized using NTA, showing that the majority of the EVs were in the size range of exosomes with a peak of around 100 nm (Supp. Figure 13B). Characterization of isolated EVs from the respective cell source confirmed surface expression of both common EV markers ALIX and TSG101, absence of the apoptotic body marker calnexin, and co-expression of both decoy proteins by WB (Supp. Figure 14). To validate the extent of engineering and to identify populations of EVs displaying both, or at least a single version, of the decoy receptors, single vesicle imaging flow cytometry^28^ was performed after labelling of EVs with anti-TNFR and anti-IL6ST antibodies (Figure 6A). As expected, we observed a heterogenous pool of engineered EVs in our analysis, where 23% of the population of engineered EVs were found to carry both decoy receptors simultaneously, whereas 37% and 40% of engineered EVs were determined to carry either TNFR1 or IL6ST respectively (Figure 6B and Supp. Figure 15A). This was further validated by immuno-gold EM, where double positive EVs could be detected (Figure 6C). Furthermore, we observed a similar trend as with single engineered EVs in the multiplex bead-based flow cytometry assay, where the surface protein profile was similar to MSC Ctrl EVs, and the decoy receptors could be detected on several sub-populations (Figure 6D and Supp. Figure 15 B-E). Decoy EVs purified from these double stable cells showed similar potency to decoy both cytokines in the *in vitro* cell assays as compared to their single decoy EV counterparts (Figure 6E-F).

**Figure 6.**
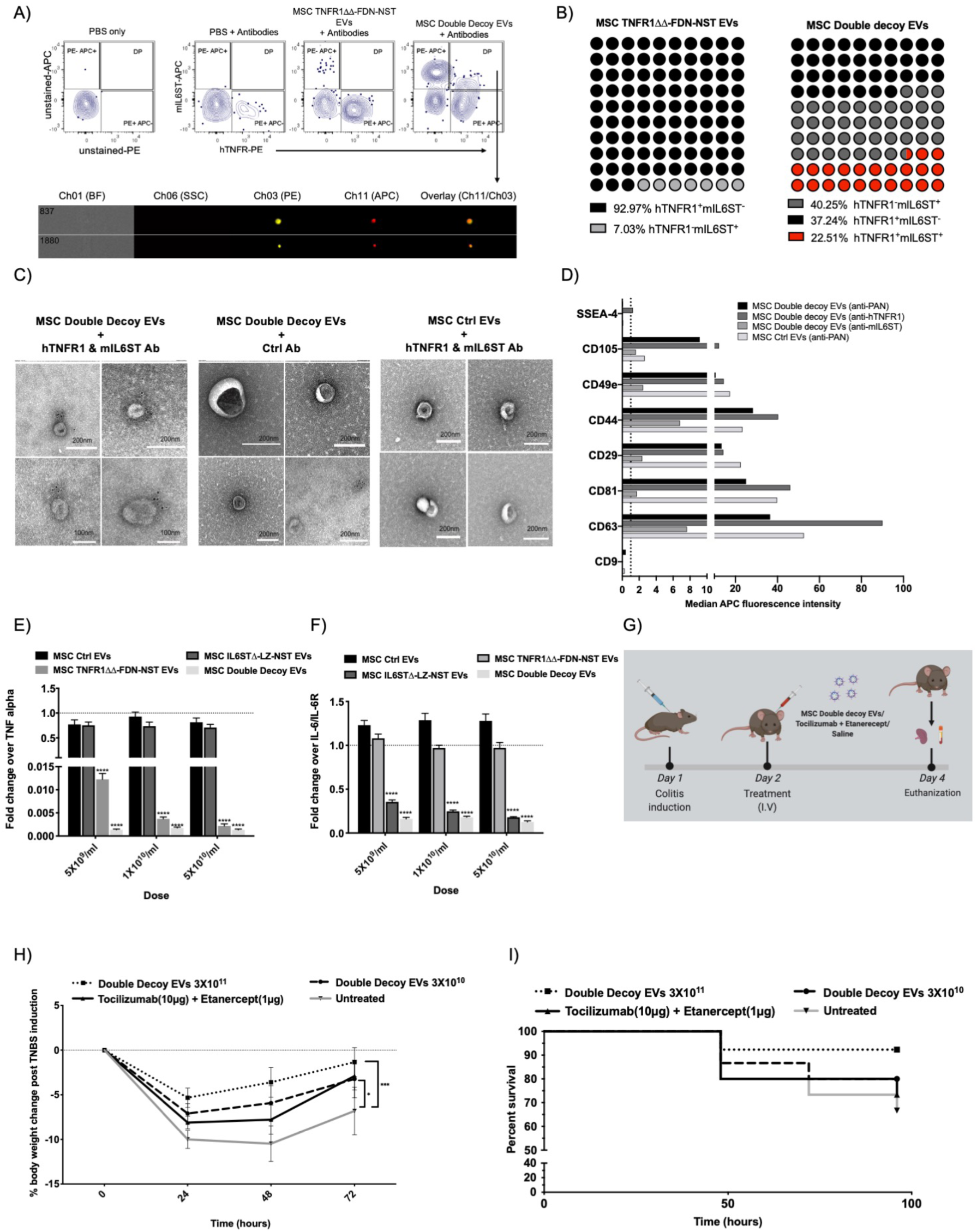
Dual functionalized Engineered EVs protect against intestinal inflammation. **A)** Imaging flow cytometry analysis (IFCM) with dot plots and example event images in the double positive (DP) gate of MSC TNFR1ΔΔ-FDN-NST and MSC double decoy EVs stained with mIL6ST APC conjugated and hTNFR1 PE conjugated antibody. PBS + antibodies were used for background adjustment and for determining the gating strategy. **B)** Percentage of detected events positive for either hTNFR1 or mIL6ST or both in Imaging flow cytometry analysis of MSC TNFR1ΔΔ-FDN-NST and MSC double decoy EVs stained with mIL6ST APC conjugated and hTNFR1 PE conjugated antibody. Percentage values determined from objects/ml in different gates. **C)** Transmission electron microscopy of double decoy EVs with nanogold labelled antibody staining of hTNFR1 (10 nm) and mIL6ST (5 nm). **D)** Multiplex EV surface characterization of PAN (CD63, CD81, and CD9) positive, hTNFR1 positive and mIL6ST positive population in MSC double decoy EVs and MSC Ctrl EVs. Data represented as background corrected median APC fluorescence intensity determined by flow cytometry of EVs bound different capture beads and upon using APC labelled detection antibody. **E)** Engineered EVs displaying TNFR1 purified from MSC cells stably expressing either the optimized TNFR1ΔΔ-FDN-NST display construct or TNFR1ΔΔ-FDN-NST and IL6STΔ-LZ-NST construct, evaluated for TNFα decoy in an *in vitro* cell assay responsive to TNFα induced NF-κB activation. EVs purified from MSC stably expressing either the IL6STΔ-LZ-NST display construct or Ctrl construct were used as control. Data were normalized to control cells treated with TNFα (5 ng/ml). **F)** Engineered EVs displaying IL6ST purified from MSC cells stably expressing either the optimized IL6STΔ-LZ-NST display construct or TNFR1ΔΔ-FDN-NST, evaluated for IL6/sIL6R decoy in an *in vitro* cell assay responsive to IL6/sIL6R induced STAT3 activation. EVs purified from MSC stably expressing either the TNFR1ΔΔ-FDN-NST display construct or Ctrl construct were used as control. Data were normalized to control cells treated with IL6/sIL6R (5 ng/ml). **G)** Schematic of the treatment protocol for double decoy EVs in TNBS induced colitis. **H)** Percent change in relative bodyweight to initial weight over the disease course and **I)** survival curve in mice induced with colitis by intrarectal injection of TNBS and treated I.V with either 3×10^11^ MSC double decoy EVs (*n*=13), 3×10^10^ MSC double decoy EVs (*n*=15), 10 *µ*g Tocilizumab and 1 *µ*g Etanercept (*n*=14), or saline (n=15) 24 hours post disease induction. **E, F**, Error bars, s.d. (*n*=3). **** *P* < 0.0001 statistical significance calculated by two-way ANOVA with Dunnett’s post-test compared with response of Ctrl EVs at the respective dose. **H**, Error bars, SEM. **** *P* < 0.0001, *** *P* < 0.001, ** *P* < 0.01, * *P*< 0.05 statistical significance calculated by two-way ANOVA with Dunnett’s post-test compared with response of Saline treated animal.

The unique ability of EVs to achieve body-wide distribution, including hard-to-reach tissues, makes them a versatile delivery vector. We and others have previously shown that upon systemic administration, EVs distribute to the gastro-intestinal tract and deliver therapeutic cargo^4,29^. Therefore, in order to validate the therapeutic effect of double decoy EVs in inhibiting intestinal inflammation *in vivo*, we used a chemically induced (TNBS) mouse model, which mimics inflammatory bowls disease (IBD). A single injection of double decoy EVs 24 hours post symptom onset, showed a significant dose-dependent reduction of weight loss at 96 hours (−1.34% for the highest dose (3×10^11^) and −3.2% for the lowest dose (3×10^10^)) compared to untreated mice (−6.8%) (Figure 6E). Importantly, the survival was also improved by double decoy EV treatment (92.3% survival) compared to untreated mice (66.7% survival) (Figure 6F). Furthermore, double decoy EVs outperformed equivalent doses (in terms of IC50 of mouse counterparts) of clinically used soluble TNFR1 protein (Etanercept) and anti-IL6R (Tocilizumab) in combination, with improved survival (92.3% for double decoy highest dose, 80.0% for the lowest dose vs 73.3% for 10 *µ*g Etanercept + 1 *µ*g Tocilizumab) and improved weight gain (1.8% weight for double decoy, highest dose, 0.8% for the lowest dose vs 0.2% for Etanercept + Tocilizumab) (Supp. Figure 16A). We also observed similar protective effect of double decoy EVs in a separate experiment with TNBS-induced colitis with improved survival (50% at 120 h) and weight change (+1.1%) compared to PBS treatment (20% survival, −1.3% weight) (Supp. Figure 16B-C). Collectively, this again confirmed the therapeutic potential of decoy EVs in inflammatory settings and further demonstrated the versatility of the engineering platform with the possibility of displaying multiple therapeutic receptors with improved efficacy.

## Discussion

With growing evidence on the critical role of EVs in a multitude of physiological processes, the potential of using EVs as a therapeutic modality has been increasingly explored. Our group and others have used various engineering strategies to achieve efficient loading of therapeutic cargos, such as nucleic acids and proteins, into EVs as well as to decorate them with various targeting ligands^1,2,5,30–33^. The approaches developed thus far rely on multi-modular engineering strategies, where a set of several proteins are used for achieving drug loading and imparting targeting moieties. Recent work by our group shows that the majority of these proteins, especially Lamp2b and CD63, fail to co-localize in the same vesicle subpopulation upon overexpression in producer cells^31^. Furthermore, Lamp2b, one of the most widely used EV engineering proteins for displaying targeting peptides, labeled only a fraction of EVs in our hands^31^. As a result of EV heterogeneity, there is a risk that for multi-modular strategies that the various modules (e.g. therapeutic and targeting) are distributed between mutually exclusive EV subpopulations, thereby negatively affecting the therapeutic efficacy of the EVs.

Here, we explored various EV engineering approaches for devising an efficient strategy to surface display therapeutic proteins. To assess the efficacy of the surface decoration and as a therapeutic application, we used the cytokine receptors TNFR1 or IL6ST, lacking their respective signaling domains. These engineered EVs were able inhibit TNFα or IL6/sIL6R complexes and hence decrease the activation of NF-κB and STAT3, respectively. This strategy allowed us to decoy pro-inflammatory cytokines in order to further treat inflammatory disorders models.

The focus of this study was to optimize the loading of the decoy receptors onto the surface of EVs by genetically modifying their producer cells. In the initial screenings, we observed efficient loading of decoy receptors onto EVs with the N-terminal fragment of Syntenin. Syntenin is a cytoplasmic adaptor of Syndecan proteoglycans and aids in the interaction of Syndecan to ALIX, a key component of the ESCRT machinery, which induces membrane budding and abscission^34^. To further enhance the efficiency of the decoy EVs, we introduced oligomerization, dimerization and trimerization domains for IL6ST and TNFR1, respectively. These were hypothesized to increase receptor decoy EV potency in two ways. First, oligomerization of exosomal sorting domains enhances the active shuttling of the cargo into EVs^35^ and second, it mimics the natural state of the receptors during ligand binding on the cell surface^12,13^. This addition of an oligomerization domain increased the inhibitory activity of TNFR1 decoy EVs by 10-fold and hence showed the importance of rigorous engineering to increase the efficacy of the EV drug, hence resulting into lower effective dosing of the treatment, thereby reducing manufacturing and production costs of this platform^36^.

To determine the therapeutic utility of these engineered EVs to suppress inflammation, we assessed the efficacy in three different inflammation models *in vivo*; LPS induced systemic inflammation, EAE, and TNBS induced colitis, which mimic sepsis, MS and IBD, respectively. In the preclinical model of sepsis, decoy EV treatments led to higher survival rate, thus indicating a dampening of the cytokine storm associated with systemic inflammation. We also observed similar effects in the preclinical model of MS, whereby systemic delivery of IL6STΔ-LZ-NST and TNFR1ΔΔ-FDN-NST decoy EVs reduced the clinical score in animals induced with EAE. Further, we demonstrated the applicability of this engineering strategy for display of other protein biologics, by simply replacing the cytokine binding region of IL6ST in IL6STΔ-LZ-NST with an alphabody against IL23. Therapeutic targeting of IL23 with IL23 decoy EVs in EAE effectively inhibited the CNS inflammation, in both prophylactic and therapeutic treatment settings. The flexibility of this platform thus allows for therapeutic applications spanning beyond decoying cytokines and could be adapted to decoy other deleterious signals, or for the display of other therapeutic and/or targeting moieties on the EV surface.

To explore the potential of displaying multiple therapeutic moieties on EVs, the same engineering strategy was used to generate double engineered EVs, carrying both TNFR1ΔΔ-FDN-NST and IL6STΔ-LZ-NST decoy receptors. These double decoy EVs showed similar efficacy as compared to single decoy EVs in *in vitro* assays for both cytokines assessed one by one, which shows that the loading of both receptors was efficient despite the similarity in engineering approach. Importantly, this points to the fact that the overexpression of the receptors does not reach the limit of the Syntenin dependent loading machinery. Furthermore, we also identified the TfR derived endosomal sorting domain to be equally efficacious compared to N terminus Syntenin fragment for IL6ST display. Although beyond the scope of this study, future developments could include conjugation of TfR domains beside Syntenin to achieve display of two or more different biologics on EV surface. However, the co-localization of these two-sorting domains into the same EV population must first be confirmed. Importantly, in the preclinical model of IBD, mice treated with double decoy EVs show improved survival and improvement in clinical symptoms as compared to a combination of clinically approved biologics against these cytokines, underpinning the benefit of decoying both cytokines simultaneously. Overall, these engineered EVs improved survival, reduced weight loss, improved clinical symptoms and down-regulated cytokine levels in the preclinical inflammation models.

Although it has been shown by us and others that exogenous EV circulation half-life in plasma is less than 5 minutes^4,37^, the precise mechanism of the therapeutic action of decoy EVs still remains to be elucidated. Importantly, EV pharmacokinetics studies have typically been performed in wild-type mice and EV biodistribution may vary in a disease relevant model. This fact was clearly highlighted in a recent study by Perets et al., where MSC EVs upon administration in various neurodegenerative models in mice, showed a completely different *in vivo* pharmacokinetics profile and primarily, with EVs accumulating in pathological CNS regions for longer durations as compared to in a healthy mouse^38^. Based on this, we speculate that cytokine decoy EVs primarily could accumulate in pro-inflammatory microenvironments upon administration. This phenomenon may explain why decoy EVs show better efficacy *in vivo* as compared to clinically used biologics.

It was recently described that EVs can be a part of an innate immune response in humans where EVs decoy bacterial toxins using ADAM10 decorated EVs^39^. Furthermore, a similar mechanism is also utilized by CD4+ T cells to decoy HIV viruses by secreting EVs enriched in CD4^40^. These studies further strengthen the therapeutic applicability of these engineered EVs for decoying toxins, soluble proteins or viruses for various clinical applications. We firmly believe that this strategy can easily be adapted for decoying e.g. viruses, merely by displaying the human proteins which serve as an entry point for the viruses. For instance, the cytokine binding parts could be replaced with human ACE2 proteolytic domain to decoy SARS-CoV-2 and prevent viral uptake into the cell. This effect can be complemented with cytokine decoy receptors to reduce the overactive immune response.

In conclusion, the platform described here has the potential to be implemented in several EV engineering strategies for displaying targeting ligands, decoy receptors, single chain antibodies, as well as other therapeutic modalities. By further modifying these designs, luminal therapeutic cargo loading and display of targeting moiety can be achieved simultaneously, hence addressing the limitation of engineering applications imparted by the heterogeneity of released EVs. By combining protein therapeutics and a natural delivery vehicle that can overcome tissue barriers, engineered EVs have great potential to be the next generation biotherapeutics.

## Supporting information

Supplementary Informatiomn

## Acknowledgments

We would like to acknowledge Bavo Vanneste and Griet Van Imschoot for their indispensable assistance. AG is an International Society for Advancement of Cytometry Marylou Ingram Scholar 2019-2023.

## Materials and Methods

### Cells

Human embryonic kidney cells (HEK293T) and immortalized human bone marrow derived mesenchymal stem cells (MSCs) were grown at 37°C, 5% CO_2_ atmosphere. HEK293T cells were cultured in Dulbecco’s Modified Eagle’s Medium (DMEM) (Invitrogen), supplemented with 10% Fetal Bovine Serum (FBS) (Invitrogen), 20 mM L-Glutamine and 1% penicillin (100 U/ml) and streptomycin (100 μg/ml) (P/S) (Sigma). MSCs were cultured in Roswell Park Memorial Institute (RPMI-1640) (Invitrogen) medium supplemented with 10% FBS, 10^−6^ mol/l hydrocortisone, and 1% P/S. 48 hours prior to harvest of conditioned medium (CM) for EV isolation, the cells were washed with PBS and media was changed to OptiMem (Invitrogen). RAW264.7 cells were grown in DMEM supplemented with 10% FBS and 1× Antibiotic-Antimycotic at 37°C in 5% CO_2_.

RAW264.7 macrophages were seeded in a 24 well-plate, at a density of 80,000 cells per well. The next day, cells were treated with 100 ng/ml of lipopolysaccharide (LPS) (L-5886, Sigma), in the presence or absence of EVs. The supernatant was collected 6 hours and 24 hours after treatment, and TNFα levels were evaluated by ELISA (BioLegend, San Diego, CA), following the manufacturer’s protocol.

### Reporter cell lines

NF-κB reporter (Luc)-HEK293 cells (BPS Bioscience, catalogue no. 60650) were cultured and used as proposed by the manufacturer. 30,000 cells per well were seeded to a 96-well plate with culture medium (DMEM + 10% FBS + 1% P/S) and incubated at 37°C with 5% CO_2_. After 24 hours, the cells were treated with or without EVs, and with hTNFα (5 ng/ml, NordicBiosite) in 50 *µ*l of complete DMEM. 6 hours after treatment the cells were lysed using 0.1% Triton X-100 in PBS (Sigma) and mixed with D-Luciferin assay (Promega) prior to luminescence measurement by Luminometer (Promega) following the manufacturer’s instructions.

HEK-Blue IL6 Cells (Invivogen, catalogue no. hkb-hil6) were cultured and used as proposed by the manufacturer. 30,000 cells per well were seeded in a 96-well plate with culture medium (DMEM + 10% FBS + 1% P/S) and incubated at 37°C with 5% CO_2_. After 24 hours, the cells were treated with or without EVs and 5 ng/ml IL6 or 5 ng/ml IL6-IL6R-complex (hyperIL6), kindly provided by Prof. Stefan Rose-John (University of Kiel, Germany). 6 hours later, an amount of 20 μl supernatant was transferred to a flat bottom 96-well plate and 180 μl of QUANTI Blue (InvivoGen) added to each well. After 3 hours incubation at 37°C the SEAP levels were quantified using a spectrophotometer (SpectraMax) at 620-655 nm.

### Plasmid constructs and cloning

For the TNFR1 display constructs, cDNA was amplified by PCR and cloned downstream of CMV promoter into a pEGFP-C1 vector backbone using NheI and BamHI. For IL6ST display constructs, codon optimized designs were synthesized (Gen9) and cloned downstream of CAG promoter into a pLEX vector backbone using EcoRI and NotI. The different constructs were assessed by transient transfection using branched polyethylenimine (PEI: total pDNA *µ*g ratio 1.5:1). Next, the complete CDS of the different display constructs was cloned into the lentiviral p2CL9IPwo5 backbone downstream of the SFFV promoter using EcoRI and NotI, and upstream of an internal ribosomal entry site-Puromycin or Neomycin resistance cDNA cassette. All expression cassettes were confirmed by sequencing and the sequences are listed in Supplementary table1.

### Production of lentiviral vectors and stable-cell lines

Lentiviral supernatants were produced as described previously^25^. In brief, HEK293T cells were co-transfected with p2CL9IPw5 plasmids containing CD63 fused to luminescent proteins, the helper plasmid pCD/NL-BH, and the human codon-optimized foamy virus envelope plasmid pcoPE using the transfection reagent JetPEI (Polyplus, Illkrich Cedex). 16 hours post transfection gene expression from the human CMV immediate-early gene enhancer/promoter was induced with 10 mM sodium butyrate (Sigma-Aldrich) for 6 hours before fresh media was added to the cells, and the supernatant was collected 22 hours later. Viral particles were pelleted at 25,000 × *g* for 90 min at 4°C. The supernatant was discarded, and the pellet was re-suspended in 2 ml of Iscove’s Modified Dulbecco’s Media supplemented with 20% FBS and 1% P/S. Aliquots were stored at −80°C until usage. To generate stable cell lines, HEK293T cells or MSC cells were transduced by overnight exposure to virus stocks and passaged at least five times under puromycin selection (Sigma; 6 *µ*g/ml)

### EV isolation

EV isolation was based on the recently optimized isolation techniques utilized in our group and described in a recent publication^41^. Briefly, conditioned media (CM) was harvested and spun first at 500 *g* for 5 minutes to remove cells, followed by 2,000 *g* for 10 minutes to remove cell debris and thereafter filtrated through an 0.22 μm filter to remove any larger particles. The CM was then run through a hollow fiber filter (D06-E300-05-N, MIDIKROS 65CM 300K MPES 0.5MM, Spectrum Laboratories) using a tangential flow filtration (TFF) system (KR2i TFF System, Spectrum Laboratories) at a flow rate of 100 ml/min (transmembrane pressure at 3.0 psi and shear rate at 3700 sec^-1^) and concentrated down to approx. 40-50 ml after diafiltration of PBS. The pre-concentrated CM was subsequently loaded onto BE-SEC columns (HiScreen Capto Core 700 column, GE Healthcare Life Sciences) and connected to an ÄKTAprime plus or ÄKTA Pure 25 chromatography system (GE Healthcare Life Sciences). Flow rate settings for column equilibration, sample loading and column cleaning in place (CIP) procedure were chosen according to the manufacturer’s instructions. The EV sample was collected according to the 280 nm UV absorbance chromatogram and concentrated using an Amicon Ultra-15 10 kDa molecular weight cut-off spin-filter (Millipore).

### Nanoparticle tracking analysis

Nanoparticle tracking analysis (NTA) was performed with a NS500 nanoparticle analyzer (NanoSight, United Kingdom) to measure the size distribution of EVs. NTA is based on the motion of nanometer-sized particles (Brownian motion) and commonly used for quantifying the concentration and size distribution of submicron-sized particles. For all our recordings, we used a camera level of 10-13 and automatic functions for all post-acquisition settings except for the detection threshold which we fixed at 6-7. Samples were diluted in PBS between 1:500 to 1:5,000 to achieve a particle count of between 2 × 10^8^ and 2 × 10^9^ per ml. The camera focus was adjusted to make the particles appear as sharp dots. Using the *script control* function, five 30 seconds videos for each sample were recorded, incorporating a sample advance and a 5 seconds delay between each recording.

### Western blot

Samples were treated with RIPA buffer and vortexed every 5 minutes for 30 minutes to lyse the EVs, subsequently the sample was spun at 12,000 *g* for 12 minutes to remove any lipids and the supernatant was collected. 30 μl of lysed sample was mixed with a sample buffer containing 0.5 M ditiothreitol (DTT), 0.4 M sodium carbonate (Na_2_CO_3_), 8% SDS and 10% glycerol, and heated at 65 °C for 5 minutes. Samples were then loaded on a NuPAGE® Novex® 4-12% Bis-Tris Gel and ran at 120 V in MOPES running buffer (Invitrogen). The proteins on the gel were transferred to an iBlot nitrocellulose membrane (Invitrogen) for 7 minutes with the iBlot system. The membranes were blocked with Odyssey blocking buffer (LiCor) diluted 1:1 in PBS for 60 minutes at room temperature with gentle shaking.

After the blocking step, the membrane was incubated with freshly prepared primary antibody solution (1:1,000 dilution for anti-Alix [ab117600, Abcam], anti-Tsg101 [ab30871, Abcam], anti-Calnexin [ab22595, Abcam], anti-His [34660, Qiagen], anti-hTNFR [ab19139, Abcam,], anti-mGp130 [R&D, #AF468] and 1:200 dilution for anti-CD81 [sc-9158, Santa Cruz]) overnight at 4°C. Membranes were washed four times, 5 minutes each using washing buffer (TBS-T 0.1%) with gentle shaking before adding the secondary antibody solution (anti-mouse IgG DyLight-800 at 1:15,000 dilution if detecting Alix or His and anti-rabbit IgG DyLight-800 at 1:15,000 dilution if detecting Calnexin or TSG101) and incubated for 1 hour at room temperature. After the secondary antibody incubation, membranes were washed four times, 5 minutes each and visualized by scanning both 700 nm and 800 nm channels on the LI-COR Odyssey CLx infrared imaging system.

### Transmission Electron Microscopy

Purified TNFR1ΔΔ-FDN-NST EVs or double decoy EVs were incubated with 1 *µ*l of 1% BSA diluted in PBS, for 5 minutes. 2 *µ*l of primary antibodies (1 mg/ml, anti-hTNFR, Abcam, ab19139) were added and incubated for 45 minutes. For the immuno-gold labeling, 2 *µ*l of protein A conjugated 10 nm gold nanoparticles (BBI Solutions) were added and incubated for 45 minutes.

Purified IL6STΔ-LZ-NST EVs or double decoy EVs were incubated with 1 *µ*l of 1% Rabbit Serum (Sigma) diluted in PBS, for 5 minutes. 2 *µ*l of primary antibody (0.2 mg/ml, anti-mGp130 from R&D, #AF468) were added and incubated for 45 minutes. For the immuno-gold labeling, 2 *µ*l of rabbit anti-goat conjugated 5 nm gold nanoparticles (BBI Solutions) were added and incubated for 45 minutes.

Finally, 3 *µ*l of labeled EVs were added onto glow-discharged formvar-carbon type B coated electron microscopy grids (Ted Pella Inc) for 3 minutes. The grid was dried with filter paper, washed twice with distilled water and blotted dry with filter paper. After the wash, the grid was stained with 2% uranyl acetate in double distilled H_2_O (Sigma) for 10 seconds and filter paper dried. The grid was air-dried and visualized on a transmission electron microscope (Tencai 10).

### Flow Cytometry

Surface expression of decoy constructs on engineered MSC lines was assessed by using either APC-conjugated rat-anti-mouse gp130 (IL6ST) antibodies (clone FAB4681A, R&D Systems) or AlexaFluor647-conjugated mouse-anti-human CD120a (TNFR1) antibodies (clone H398, Bio-Rad). DAPI was used for dead cell exclusion.

### Multiplex bead-based flow cytometry analysis

Multiplex bead-based flow cytometry analysis (MACSPlex Exosome Kit, human, Miltenyi Biotec) was implemented to characterize general surface protein composition of decoy EVs and specific surface proteins co-expressed on engineered decoy receptor EVs. Assays were performed based on an optimized protocol described previously^25^. In brief, EVs were used at an input dose of 1×10^9^ NTA-based particles per assay, diluted with MACSPlex buffer to a total volume of 120 *µ*l and incubated with 15 *µ*l of MACSPlex Exosome Capture Beads overnight in wells of a pre-wet and drained MACSPlex 96 well 0.22 *µ*m filter plate at 450 *g* at room temperature. Beads were washed with 200 *µ*l MACSPlex buffer and the liquid was removed applying vacuum (Sigma-Aldrich, Supelco PlatePrep; −100 mBar). For counterstaining of captured EVs, either a mixture of APC-conjugated anti-CD9, anti-CD63 and anti-CD81 detection antibodies (supplied in the MACSPlex kit, 5 *µ*l each) or anti-decoy receptor antibodies (AlexaFluor647-labelled anti-human TNFR1, Bio-Rad, cat #MCA1340A647, clone H398, lot 0410; or APC-labelled anti-mouse gp130, R&D Systems, cat #FAB4681A, clone 125623, lot AAOK0114071; 200 ng, respectively) were added to each well in a total volume of 135 *µ*l and the plate was incubated at 450 *g* for 1 hours at room temperature. Next, the samples were washed twice, resuspended in MACSPlex buffer and analyzed by flow cytometry with a MACSQuant Analyzer 10 flow cytometer (Miltenyi Biotec). FlowJo software (version 10.6.2, FlowJo, LLC) was used to analyze flow cytometric data. Median fluorescence intensities (MFI) for all 39 capture bead subsets were background-corrected by subtracting respective MFI values from matched non-EV containing buffer controls (buffer + capture beads + antibodies) that were treated exactly like sEV-containing samples (buffer + capture beads + EVs + antibodies).

### Single EV analysis by Imaging Flow Cytometry

TNFR-decoy EVs and double decoy EVs were analyzed by single EV Imaging Flow Cytometry (IFCM) to confirm decoy receptor co-expression on a portion of engineered EVs. Isolated EVs were stained with PE-labelled anti-human TNFR1 (Bio-Rad, cat #MCA1340PE, clone H398, lot 0407) and APC-labelled anti-mouse gp130 antibodies (R&D Systems, cat #FAB4681A, clone 125623, lot AAOK0114071; final concentration during staining 10 nM) at a concentration of 5×10^9^ NTA-based particles in 60 *µ*l total volume for 1 hour at room temperature. Samples (and buffer controls without EVs, i.e. PBS + antibodies) were diluted 200-fold in PBS post staining and analyzed using an ImageStreamX MkII instrument (ISX; Amnis/Luminex) equipped with 5 lasers (70 mW 375 nm, 100 mW 488 nm, 200 mW 561 nm, 150 mW 642 nm, 70 mW 785 nm [SSC]) with a protocol and masking setting described and optimized previously^28,42^. Fluorescence parameters recorded and analyzed in this study comprise PE signals (MC_Or_NMC_Channel_03) and APC/AlexaFluor 647 (MC_Or_NMC_Channel_11). All analyses were performed by using the 60× objective and deactivated Remove Beads option. All lasers were set to maximum powers, and all data was acquired with a 7 μm core size and low flow rate (∼0.38 μL/min). Data was recorded for 5 min and pre-gated on SSC (low) events as described previously. Dulbecco’s PBS pH 7.4 (Gibco) was used as sheath fluid and for all dilution steps. Data was analyzed with optimized masking settings and by excluding coincidence events as described before using Amnis IDEAS software (version 6.2.187.0) and FlowJo v. 10.6.2 (FlowJo, LLC).

### Reverse Transcription-quantitative PCR (RT-qPCR)

Total RNA was isolated using TRIzol reagent (Invitrogen) and Aureum Total RNA Isolation Mini Kit (Bio-Rad), according to manufacturer’s instructions. cDNA synthesis was performed using iScript cDNA synthesis kit (Bio-Rad Laboratories), according to manufacturer’s instructions. RT-qPCR was performed with the Light Cycler 480 system (Roche) using Sensifast Bioline Mix (Bio-Line). Expression levels in the spinal cord were normalized to the expression of the two most stable housekeeping genes, which were determined using geNorm^43^: ubiquitin-C (*Ubc*) and hypoxanthine-guanine phosphoribosyltransferase (*Hprt*).

Primer sequences:

Ubc (Fw 5′-AGGTCAAACAGGAAGACAGACGTA-

3’, Rev 5’-TCACACCCAAGAACAAGCACA-3’),

Hprt (Fw 5’-AGTGTTGGATACAGGCCAGAC-3’, Rev

5’-CGTGATTCAAATCCCTGAAGT-3’),

Il6 (Fw 5’-TAGTCCTTCCTACCCCAATTTCC-3’, Rev

5’-TTGGTCCTTAGCCACTCCTTC-3’),

Tnf (Fw 5’-ACCCTGGTATGAGCCCATATAC-3’, Rev

5’-ACACCCATTCCCTTCACAGAG-3’),

Il17a (Fw 5’-TTTAACTCCCTTGGCGCAAAA-3’, Rev

5’-CTTTCCCTCCGCATTGACAC-3’),

Cxcl1 (Fw 5’-CTGGGATTCACCTCAAGAACATC-3’, Rev

5’-CAGGGTCAAGGCAAGCCTC-3’).

### Animal experiments

#### Systemic inflammation model

Systemic inflammation was induced using Lipopolysaccharide (LPS) as described by others. 20 g (±5 g) female C57BL/6 mice were injected intraperitoneally (I.P) with LPS (L-5886, Sigma). EVs were I.V injected via the tail vein subsequent to LPS induction and the animals were observed and weighed daily after induction. The animal experiments were approved by The Swedish Local Board for Laboratory Animals.

#### Experimental autoimmune encephalitis (EAE) model

EAE was induced as described previously^19^. 20 g (±5 g) female C57BL/6 mice were immunized by subcutaneous S.C injection of 100 *µ*l of the MOG35-55-CFA emulsion subcutaneously, distributed to 3 different locations. I.P injections of 400 ng pertussis toxin were given on the day of and two days following immunization to induce disease. Mice were subsequently monitored for change in body weight and assessed using EAE-scoring, see Table 1. EVs were injected either S.C on day 7, 9 and 13 or given as single I.V injection. The animal experiments were approved by The Swedish Local Board for Laboratory Animals.

#### TNBS induced colitis model

Trinitrobenzene sulfonic acid (TNBS) induced Colitis was induced as described previously^44^. 20 g (±5 g) female BALB/c mice were pre-sensitized with peritoneum skin application of 60 *µ*l 5% TNBS + 90 *µ*l acetone-olive oil (4:1) mix per mouse. One week later, Colitis was induced by intra-rectal administration of 30 *µ*l TNBS + 42.1 *µ*l 95% ethanol + 27.9 *µ*l H_2_O per mouse. Mice were subsequently monitored for change in body weight.

All animal experiments were performed in accordance with the ethical permission and designed to minimize the suffering and pain of the animals.

#### Statistical analysis

Statistical analyses of the data were performed using Prism 6.0 (GraphPad Software Inc.) by one-way ANOVA or two-way ANOVA for all *P*-values. All results are expressed as mean ±SEM. All graphs were made in Prism 6.0 (GraphPad Software Inc.).

## References

(1) Armstrong, J. P. K.; Holme, M. N.; Stevens, M. M. Re-Engineering Extracellular Vesicles as Smart Nanoscale Therapeutics. ACS Nano 2017, 11 (1), 69–83. https://doi.org/10.1021/acsnano.6b07607.

(2) Alvarez-Erviti, L.; Seow, Y.; Yin, H.; Betts, C.; Lakhal, S.; Wood, M. J. A. Delivery of SiRNA to the Mouse Brain by Systemic Injection of Targeted Exosomes. Nat. Biotechnol. 2011, 29 (4), 341–345. https://doi.org/10.1038/nbt.1807.

(3) El-Andaloussi, S.; Lee, Y.; Lakhal-Littleton, S.; Li, J.; Seow, Y.; Gardiner, C.; Alvarez-Erviti, L.; Sargent, I. L.; Wood, M. J. A. Exosome-Mediated Delivery of SiRNA in Vitro and in Vivo. Nat. Protoc. 2012, 7 (12), 2112–2126. https://doi.org/10.1038/nprot.2012.131.

(4) Wiklander, O. P. B.; Nordin, J. Z.; O’Loughlin, A.; Gustafsson, Y.; Corso, G.; Mäger, I.; Vader, P.; Lee, Y.; Sork, H.; Seow, Y.; et al. Extracellular Vesicle in Vivo Biodistribution Is Determined by Cell Source, Route of Administration and Targeting. J. Extracell. Vesicles 2015. https://doi.org/10.3402/jev.v4.26316.

(5) Cooper, J. M.; Wiklander, P. B. O.; Nordin, J. Z.; Al-Shawi, R.; Wood, M. J.; Vithlani, M.; Schapira, A. H. V.; Simons, J. P.; El-Andaloussi, S.; Alvarez-Erviti, L. Systemic Exosomal SiRNA Delivery Reduced Alpha-Synuclein Aggregates in Brains of Transgenic Mice. Mov. Disord. 2014, 29 (12), 1476–1485. https://doi.org/10.1002/mds.25978.

(6) Jeppesen, D. K.; Fenix, A. M.; Franklin, J. L.; Higginbotham, J. N.; Zhang, Q.; Zimmerman, L. J.; Liebler, D. C.; Ping, J.; Liu, Q.; Evans, R.; et al. Reassessment of Exosome Composition. Cell 2019, 177 (2), 428-445.e18. https://doi.org/10.1016/j.cell.2019.02.029.

(7) Willms, E.; Johansson, H. J.; Mäger, I.; Lee, Y.; Blomberg, K. E. M.; Sadik, M.; Alaarg, A.; Smith, C. I. E.; Lehtiö, J.; El Andaloussi, S.; et al. Cells Release Subpopulations of Exosomes with Distinct Molecular and Biological Properties. Sci. Rep. 2016, 6. https://doi.org/10.1038/srep22519.

(8) Vagner, T.; Chin, A.; Mariscal, J.; Bannykh, S.; Engman, D. M.; Di Vizio, D. Protein Composition Reflects Extracellular Vesicle Heterogeneity. Proteomics 2019, 19 (8), 1800167. https://doi.org/10.1002/pmic.201800167.

(9) Crescitelli, R.; Lässer, C.; Jang, S. C.; Cvjetkovic, A.; Malmhäll, C.; Karimi, N.; Höög, J. L.; Johansson, I.; Fuchs, J.; Thorsell, A.; et al. Subpopulations of Extracellular Vesicles from Human Metastatic Melanoma Tissue Identified by Quantitative Proteomics after Optimized Isolation. J. Extracell. Vesicles 2020, 9 (1). https://doi.org/10.1080/20013078.2020.1722433.

(10) Kim, E. Y.; Moudgil, K. D. Immunomodulation of Autoimmune Arthritis by Pro-Inflammatory Cytokines. Cytokine. Academic Press October 1, 2017, pp 87–96. https://doi.org/10.1016/j.cyto.2017.04.012.

(11) Moudgil, K. D.; Choubey, D. Cytokines in Autoimmunity: Role in Induction, Regulation, and Treatment. Journal of Interferon and Cytokine Research. Mary Ann Liebert Inc. October 1, 2011, pp 695–703. https://doi.org/10.1089/jir.2011.0065.

(12) Garbers, C.; Heink, S.; Korn, T.; Rose-John, S. Interleukin-6: Designing Specific Therapeutics for a Complex Cytokine. Nature Reviews Drug Discovery. Nature Publishing Group June 1, 2018, pp 395–412. https://doi.org/10.1038/nrd.2018.45.

(13) Kalliolias, G. D.; Ivashkiv, L. B. TNF Biology, Pathogenic Mechanisms and Emerging Therapeutic Strategies. Nat. Rev. Rheumatol. 2016, 12 (1), 49–62. https://doi.org/10.1038/nrrheum.2015.169.

(14) Sedger, L. M.; McDermott, M. F. TNF and TNF-Receptors: From Mediators of Cell Death and Inflammation to Therapeutic Giants - Past, Present and Future. Cytokine and Growth Factor Reviews. Elsevier Ltd August 1, 2014, pp 453–472. https://doi.org/10.1016/j.cytogfr.2014.07.016.

(15) Wolf, J.; Rose-John, S.; Garbers, C. Interleukin-6 and Its Receptors: A Highly Regulated and Dynamic System. Cytokine 2014, 70 (1), 11–20. https://doi.org/10.1016/j.cyto.2014.05.024.

(16) Simpson, R. J.; Kalra, H.; Mathivanan, S. Exocarta as a Resource for Exosomal Research. J. Extracell. Vesicles 2012, 1 (1). https://doi.org/10.3402/jev.v1i0.18374.

(17) Hurwitz, S. N.; Rider, M. A.; Bundy, J. L.; Liu, X.; Singh, R. K.; Meckes, D. G. Proteomic Profiling of NCI-60 Extracellular Vesicles Uncovers Common Protein Cargo and Cancer Type-Specific Biomarkers. Oncotarget 2016, 7 (52), 86999–87015. https://doi.org/10.18632/oncotarget.13569.

(18) Sork, H.; Corso, G.; Krjutskov, K.; Johansson, H. J.; Nordin, J. Z.; Wiklander, O. P. B.; Lee, Y. X. F.; Westholm, J. O.; Lehtiö, J.; Wood, M. J. A.; et al. Heterogeneity and Interplay of the Extracellular Vesicle Small RNA Transcriptome and Proteome. Sci. Rep. 2018, 8 (1). https://doi.org/10.1038/s41598-018-28485-9.

(19) Xanthoulea, S.; Pasparakis, M.; Kousteni, S.; Brakebusch, C.; Wallach, D.; Bauer, J.; Lassmann, H.; Kollias, G. Tumor Necrosis Factor (TNF) Receptor Shedding Controls Thresholds of Innate Immune Activation That Balance Opposing TNF Functions in Infectious and Inflammatory Diseases. J. Exp. Med. 2004, 200 (3), 367–376. https://doi.org/10.1084/jem.20040435.

(20) Meier, S.; Güthe, S.; Kiefhaber, T.; Grzesiek, S. Foldon, the Natural Trimerization Domain of T4 Fibritin, Dissociates into a Monomeric A-State Form Containing a Stable β-Hairpin: Atomic Details of Trimer Dissociation and Local β-Hairpin Stability from Residual Dipolar Couplings. J. Mol. Biol. 2004, 344 (4), 1051–1069. https://doi.org/10.1016/j.jmb.2004.09.079.

(21) Ellenberger, T. E.; Brandl, C. J.; Struhl, K.; Harrison, S. C. The GCN4 Basic Region Leucine Zipper Binds DNA as a Dimer of Uninterrupted α Helices: Crystal Structure of the Protein-DNA Complex. Cell 1992, 71 (7), 1223–1237. https://doi.org/10.1016/S0092-8674(05)80070-4.

(22) Jensen, M. R.; Yabukarski, F.; Communie, G.; Condamine, E.; Mas, C.; Volchkova, V.; Tarbouriech, N.; Bourhis, J.-M.; Volchkov, V.; Blackledge, M.; et al. STRUCTURAL DESCRIPTION OF THE NIPAH VIRUS PHOSPHOPROTEIN AND ITS INTERACTION WITH STAT1. Biophys. J. 2020. https://doi.org/10.1016/j.bpj.2020.04.010.

(23) Sliepen, K.; Van Montfort, T.; Melchers, M.; Isik, G.; Sanders, R. W. Immunosilencing a Highly Immunogenic Protein Trimerization Domain. J. Biol. Chem. 2015, 290 (12), 7436–7442. https://doi.org/10.1074/jbc.M114.620534.

(24) Théry, C.; Witwer, K. W.; Aikawa, E.; Alcaraz, M. J.; Anderson, J. D.; Andriantsitohaina, R.; Antoniou, A.; Arab, T.; Archer, F.; Atkin-Smith, G. K.; et al. Minimal Information for Studies of Extracellular Vesicles 2018 (MISEV2018): A Position Statement of the International Society for Extracellular Vesicles and Update of the MISEV2014 Guidelines. J. Extracell. Vesicles 2018, 7 (1). https://doi.org/10.1080/20013078.2018.1535750.

(25) Wiklander, O. P. B.; Bostancioglu, R. B.; Welsh, J. A.; Zickler, A. M.; Murke, F.; Corso, G.; Felldin, U.; Hagey, D. W.; Evertsson, B.; Liang, X.-M.; et al. Systematic Methodological Evaluation of a Multiplex Bead-Based Flow Cytometry Assay for Detection of Extracellular Vesicle Surface Signatures. Front. Immunol. 2018, 9, 1326. https://doi.org/10.3389/fimmu.2018.01326.

(26) Desmet, J.; Verstraete, K.; Bloch, Y.; Lorent, E.; Wen, Y.; Devreese, B.; Vandenbroucke, K.; Loverix, S.; Hettmann, T.; Deroo, S.; et al. Structural Basis of IL23 Antagonism by an Alphabody Protein Scaffold. Nat. Commun. 2014, 5 (1), 1–12. https://doi.org/10.1038/ncomms6237.

(27) Lovett-Racke, A. E.; Racke, M. K. Role of IL-12/IL23 in the Pathogenesis of Multiple Sclerosis. In Neuroinflammation; Elsevier, 2018; pp 115–139. https://doi.org/10.1016/b978-0-12-811709-5.00005-3.

(28) Görgens, A.; Bremer, M.; Ferrer-Tur, R.; Murke, F.; Tertel, T.; Horn, P. A.; Thalmann, S.; Welsh, J. A.; Probst, C.; Guerin, C.; et al. Optimisation of Imaging Flow Cytometry for the Analysis of Single Extracellular Vesicles by Using Fluorescence-Tagged Vesicles as Biological Reference Material. J. Extracell. Vesicles 2019, 8 (1). https://doi.org/10.1080/20013078.2019.1587567.

(29) Reshke, R.; Taylor, J. A.; Savard, A.; Guo, H.; Rhym, L. H.; Kowalski, P. S.; Trung, M. T.; Campbell, C.; Little, W.; Anderson, D. G.; et al. Reduction of the Therapeutic Dose of Silencing RNA by Packaging It in Extracellular Vesicles via a Pre-MicroRNA Backbone. Nat. Biomed. Eng. 2020. https://doi.org/10.1038/s41551-019-0502-4.

(30) Kamerkar, S.; Lebleu, V. S.; Sugimoto, H.; Yang, S.; Ruivo, C. F.; Melo, S. A.; Lee, J. J.; Kalluri, R. Exosomes Facilitate Therapeutic Targeting of Oncogenic KRAS in Pancreatic Cancer. Nature 2017, 546 (7659), 498–503. https://doi.org/10.1038/nature22341.

(31) Corso, G.; Heusermann, W.; Trojer, D.; Görgens, A.; Steib, E.; Voshol, J.; Graff, A.; Genoud, C.; Lee, Y.; Hean, J.; et al. Systematic Characterization of Extracellular Vesicle Sorting Domains and Quantification at the Single Molecule – Single Vesicle Level by Fluorescence Correlation Spectroscopy and Single Particle Imaging. J. Extracell. Vesicles 2019, 8 (1), 1663043. https://doi.org/10.1080/20013078.2019.1663043.

(32) Gao, X.; Ran, N.; Dong, X.; Zuo, B.; Yang, R.; Zhou, Q.; Moulton, H. M.; Seow, Y.; Yin, H. F. Anchor Peptide Captures, Targets, and Loads Exosomes of Diverse Origins for Diagnostics and Therapy. Sci. Transl. Med. 2018, 10 (444). https://doi.org/10.1126/scitranslmed.aat0195.

(33) Wiklander, O. P. B.; Brennan, M.; Lötvall, J.; Breakefield, X. O.; Andaloussi, S. E. L. Advances in Therapeutic Applications of Extracellular Vesicles. Science Translational Medicine. American Association for the Advancement of Science 2019. https://doi.org/10.1126/scitranslmed.aav8521.

(34) Baietti, M. F.; Zhang, Z.; Mortier, E.; Melchior, A.; Degeest, G.; Geeraerts, A.; Ivarsson, Y.; Depoortere, F.; Coomans, C.; Vermeiren, E.; et al. Syndecan-Syntenin-ALIX Regulates the Biogenesis of Exosomes. Nat. Cell Biol. 2012, 14 (7), 677–685. https://doi.org/10.1038/ncb2502.

(35) Fang, Y.; Wu, N.; Gan, X.; Yan, W.; Morrell, J. C.; Gould, S. J. Higher-Order Oligomerization Targets Plasma Membrane Proteins and HIV Gag to Exosomes. PLoS Biol. 2007, 5 (6), e158. https://doi.org/10.1371/journal.pbio.0050158.

(36) Whitford, W.; Guterstam, P. Exosome Manufacturing Status. https://doi.org/10.4155/fmc-2018-0417.

(37) Lai, C. P.; Mardini, O.; Ericsson, M.; Prabhakar, S.; Maguire, C. A.; Chen, J. W.; Tannous, B. A.; Breakefield, X. O. Dynamic Biodistribution of Extracellular Vesicles in Vivo Using a Multimodal Imaging Reporter. ACS Nano 2014, 8 (1), 483–494. https://doi.org/10.1021/nn404945r.

(38) Perets, N.; Betzer, O.; Shapira, R.; Brenstein, S.; Angel, A.; Sadan, T.; Ashery, U.; Popovtzer, R.; Offen, D. Golden Exosomes Selectively Target Brain Pathologies in Neurodegenerative and Neurodevelopmental Disorders. Nano Lett. 2019, 19 (6), 3422–3431. https://doi.org/10.1021/acs.nanolett.8b04148.

(39) Keller, M. D.; Ching, K. L.; Liang, F. X.; Dhabaria, A.; Tam, K.; Ueberheide, B. M.; Unutmaz, D.; Torres, V. J.; Cadwell, K. Decoy Exosomes Provide Protection against Bacterial Toxins. Nature 2020, 579 (7798), 260–264. https://doi.org/10.1038/s41586-020-2066-6.

(40) de Carvalho, J. V.; de Castro, R. O.; da Silva, E. Z. M.; Silveira, P. P.; da Silva-Januário, M. E.; Arruda, E.; Jamur, M. C.; Oliver, C.; Aguiar, R. S.; daSilva, L. L. P. Nef Neutralizes the Ability of Exosomes from CD4+ T Cells to Act as Decoys during HIV-1 Infection. PLoS One 2014, 9 (11), e113691. https://doi.org/10.1371/journal.pone.0113691.

(41) Corso, G.; Mäger, I.; Lee, Y.; Görgens, A.; Bultema, J.; Giebel, B.; Wood, M. J. A.; Nordin, J. Z.; Andaloussi, S. El. Reproducible and Scalable Purification of Extracellular Vesicles Using Combined Bind-Elute and Size Exclusion Chromatography. Sci. Rep. 2017, 7 (1). https://doi.org/10.1038/s41598-017-10646-x.

(42) Tertel, T., Bremer, M., Maire, C.L., Lamszus, K., Peine, S., Jawad, R., El Andaloussi, S., Giebel, B., Ricklefs, F.L., and Gorgens, A. High-Resolution Imaging Flow Cytometry Reveals Impact of Incubation Temperature on Labelling of Extracellular Vesicles with Antibodies. Cytom. Part A 2020.

(43) Brkic, M.; Balusu, S.; Van Wonterghem, E.; Gorlé, N.; Benilova, I.; Kremer, A.; Van Hove, I.; Moons, L.; De Strooper, B.; Kanazir, S.; et al. Amyloid β Oligomers Disrupt Blood–CSF Barrier Integrity by Activating Matrix Metalloproteinases. J. Neurosci. 2015, 35 (37), 12766–12778. https://doi.org/10.1523/JNEUROSCI.0006-15.2015.

(44) Scheiffele, F.; Fuss, I. J. Induction of TNBS Colitis in Mice. In Current Protocols in Immunology; John Wiley & Sons, Inc., 2002. https://doi.org/10.1002/0471142735.im1519s49.

