## Supplementary Informatiomn for "Engineering of extracellular vesicles for display of protein biotherapeutics"

**Supplementary information**

Supplementary Figure 1

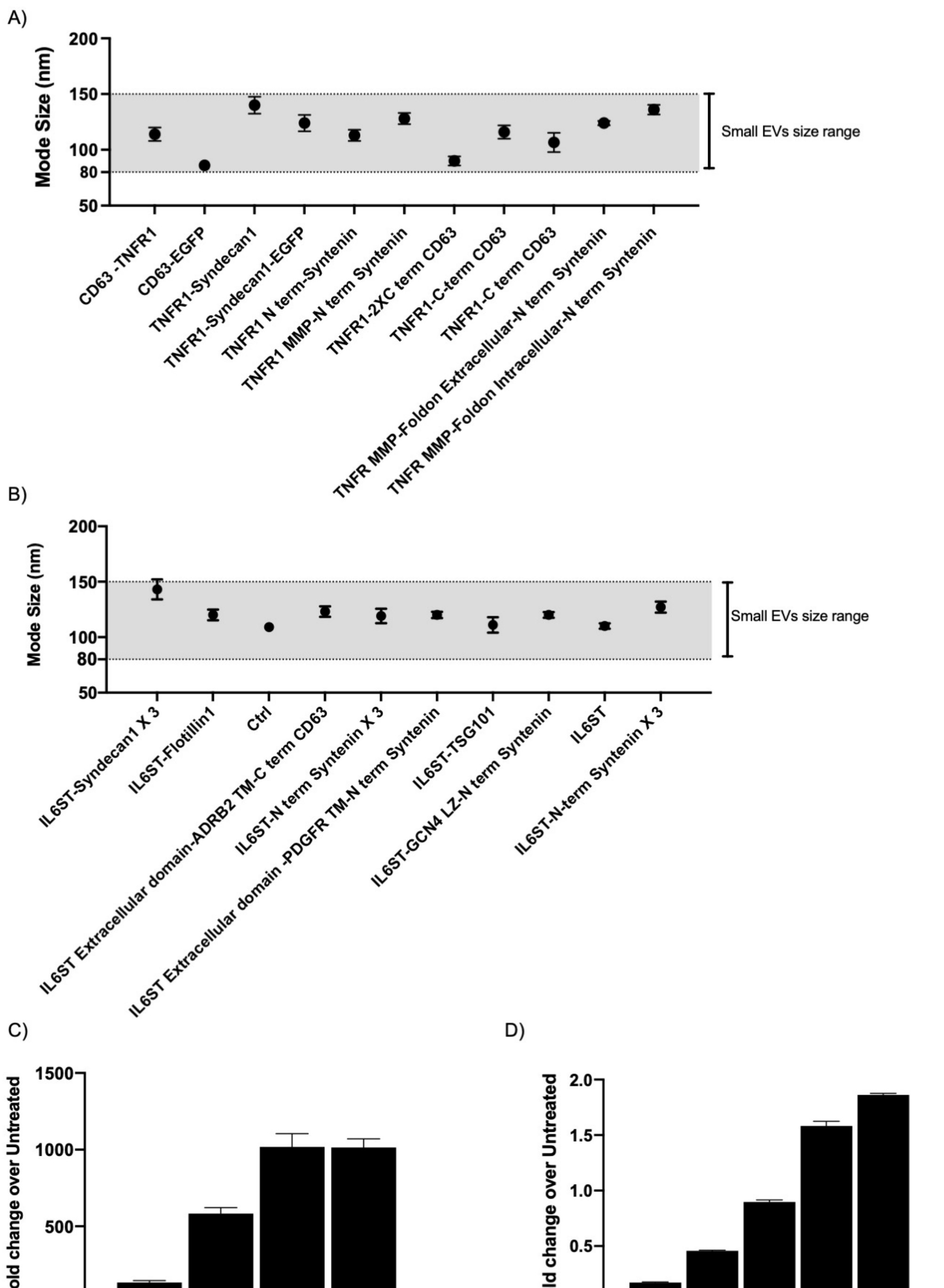


**Supplementary Figure 1. A-B)** Mode size determined by NTA of indicated engineered HEK293T EVs purified from cells transfected with various A) TNFR1 display constructs and B) IL6ST display constructs. **C)** Fold change in signal over untreated HEK293T NF-κB reporter cells, upon incubation of different doses of TNFα for 4 hours. **D)** Fold change in signal over untreated HEK293T STAT3 reporter cells, upon incubation of different doses of IL6/sIL6R for 12 hours.

#### Supplementary Figure 2

A)

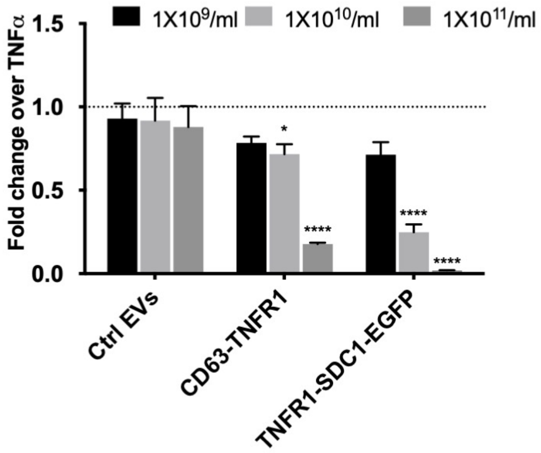


B)

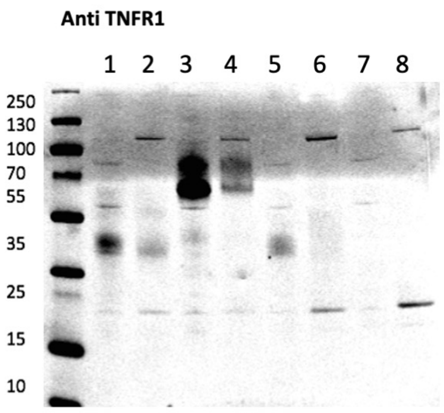


| Lane | Sample Name | Predicted M.W kDa |
|---|---|---|
| 1 | TNFR1-Syndecan1 Cells | 42 |
| 2 | TNFR1-Syndecan1 EVs | 42 |
| 3 | TNFR1-Syndecan1-EGFP Cells | 59 |
| 4 | TNFR1-Syndecan1-EGFP EVs | 59 |
| 5 | CD63-TNFR1 Cells | 40 |
| 6 | CD63-TNFR1 EVs | 40 |
| 7 | Ctrl Cells | - |
| 8 | Ctrl EVs | - |

**Supplementary Figure 2. A)** Engineered EVs displaying TNFR1 purified from HEK293T cells transfected with constructs listed in the figure legends, evaluated for TNFα decoy in an *in vitro* cell assay respondent to TNFα induced NF-ϰB activation. Data were normalized to control cells treated with TNFα (5ng/ml). Error bars, s.d. ($n$=3). **** $P < 0.0001$ statistical significance calculated by two-way ANOVA upon comparing response to Ctrl EVs at the respective dose. **B)** WB probed against human TNFR1 on engineered HEK293T EVs and their respective cells transfected with various EVs engineering designs. Predicted molecular weight (kDa) was calculated based on the amino acid sequence of the respective construct.

#### Supplementary Figure 3

A)

**Anti poly-Histidine tag**

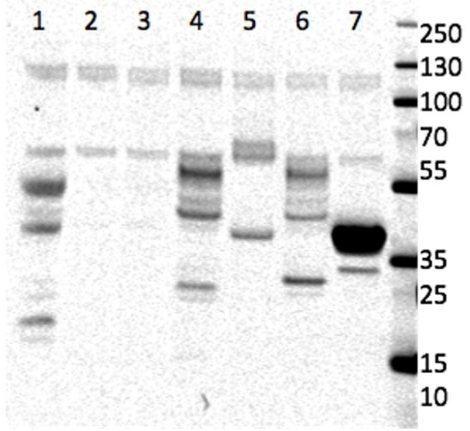


| Lane | Sample Name | Predicted M.W kDa |
|---|---|---|
| 1 | TNFR1 Cells | 38 |
| 2 | TNFR1-C term CD63 Cells | 40 |
| 3 | TNFR1-2XC term CD63 Cells | 42 |
| 4 | TNFR1-Syndecan1-EGFP Cells | 42 |
| 5 | TNFR1-Syndecan1 Cells | 59 |
| 6 | TNFR1-N term Syntenin Cells | 45 |
| 7 | Ctrl Cells | - |

B)

**Anti poly-Histidine tag**

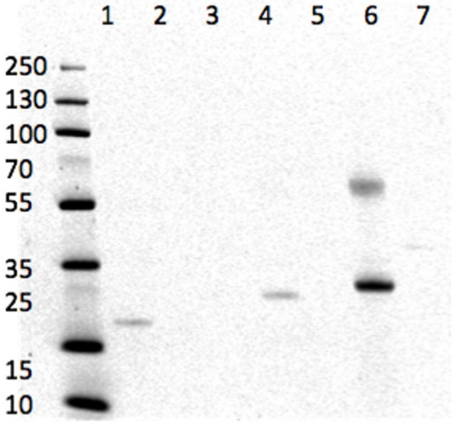


| Lane | Sample Name | Predicted M.W kDa |
|---|---|---|
| 1 | TNFR1 EVs | 38 |
| 2 | TNFR1-C term CD63 EVs | 40 |
| 3 | TNFR1-2XC term CD63 EVs | 42 |
| 4 | TNFR1-Syndecan1-EGFP EVs | 42 |
| 5 | TNFR1-Syndecan1 EVs | 59 |
| 6 | TNFR1-N term Syntenin EVs | 45 |
| 7 | Ctrl EVs | - |

**Supplementary Figure 3.** WB probed against poly Histidine tag on **A)** engineered HEK293T cells and their **B)** respective EVs. Cells were transfected with indicated genetic constructs. Predicted molecular weight (kDa) was calculated based on the amino acid sequence of the respective construct.

#### Supplementary Figure 4

A)

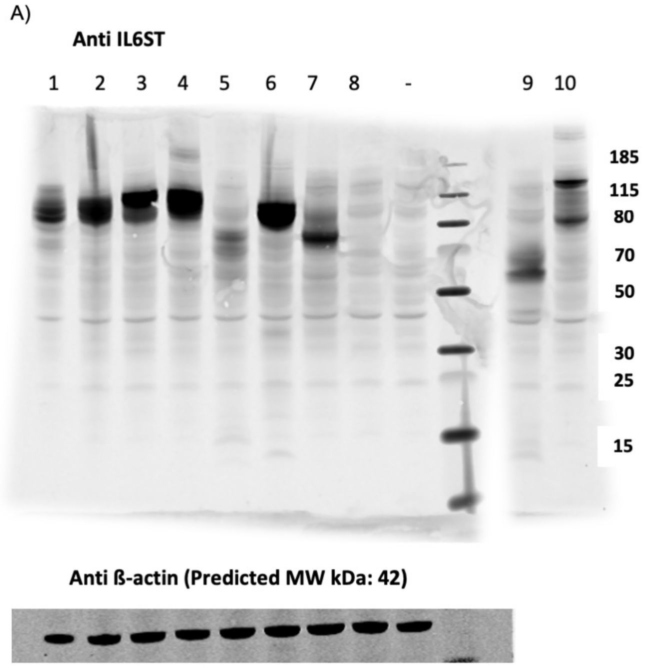


| Lane | Sample Name | Predicted M.W kDa |
|---|---|---|
| 1 | IL6ST-3X N term Syntenin Cells | 118 |
| 2 | IL6ST-N term Syntenin Cells | 91 |
| 3 | IL6ST-Syndecan1 Cells | 93 |
| 4 | IL6ST-GCN4 L.Z-N term Syntenin Cells | 94 |
| 5 | IL6ST extracellular domain-PDGFR TM-N term Syntenin Cells | 57 |
| 6 | IL6ST Cells | 77 |
| 7 | IL6ST extracellular domain-ADRB2 TM-C term CD63 Cells | 70 |
| 8 | Ctrl Cells | |
| 9 | IL6ST extracellular domain-PDGFR TM Cells | 43 |
| 10 | IL6ST-TSG101 Cells | 121 |

B)

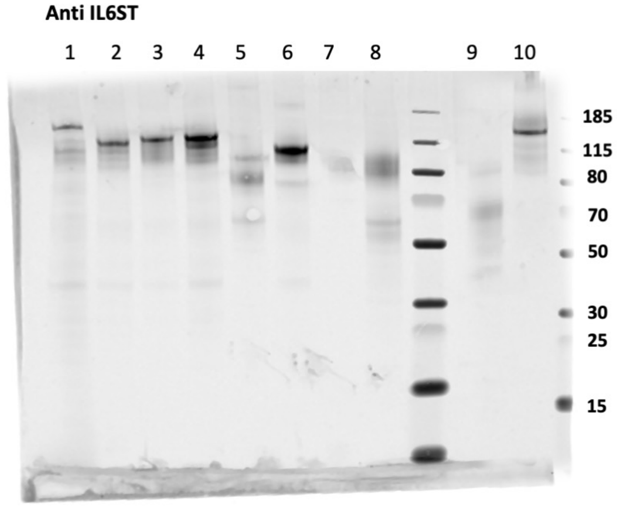


| Lane | Sample Name | Predicted M.W kDa |
|---|---|---|
| 1 | IL6ST-3X N term Syntenin EVs | 118 |
| 2 | IL6ST-N term Syntenin EVs | 91 |
| 3 | IL6ST-Syndecan1 EVs | 93 |
| 4 | IL6ST-GCN4 L.Z-N term Syntenin EVs | 94 |
| 5 | IL6ST extracellular domain-PDGFR TM-N term Syntenin EVs | 57 |
| 6 | IL6ST EVs | 77 |
| 7 | IL6ST extracellular domain-ADRB2 TM-C term CD63 EVs | 70 |
| 8 | Ctrl EVs | |
| 9 | IL6ST extracellular domain-PDGFR TM EVs | 43 |
| 10 | IL6ST-TSG101 EVs | 121 |

**Supplementary Figure 4.** WB probed against mouse IL6ST on **A)** engineered HEK293T cells and their **B)** respective EVs. Cells were transfected with indicated genetic constructs. Predicted molecular weight (kDa) values were calculated based on the amino acid sequence of the respective construct.

#### Supplementary Figure 5

A)

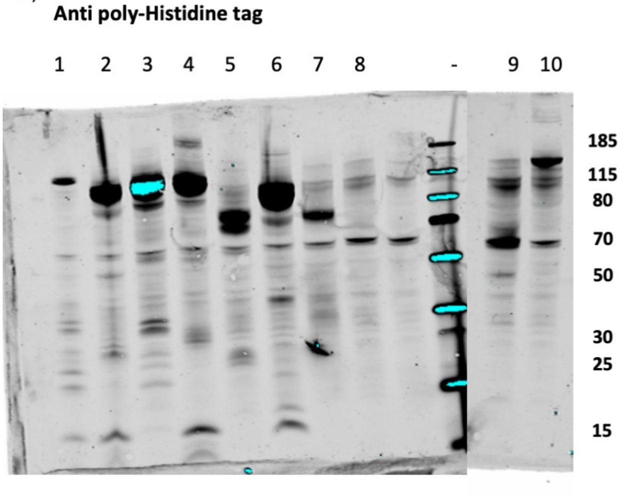


| Lane | Sample Name | Predicted M.W kDa |
|---|---|---|
| 1 | IL6ST-3X N term Syntenin Cells | 118 |
| 2 | IL6ST-N term Syntenin Cells | 91 |
| 3 | IL6ST-Syndecan1 Cells | 93 |
| 4 | IL6ST-GCN4 L.Z-N term Syntenin Cells | 94 |
| 5 | IL6ST extracellular domain-PDGFR TM-N term Syntenin Cells | 57 |
| 6 | IL6ST Cells | 77 |
| 7 | IL6ST extracellular domain-ADRB2 TM-CD63 Cells | 70 |
| 8 | Ctrl Cells | |
| 9 | IL6ST extracellular domain-PDGFR TM Cells | 43 |
| 10 | IL6ST-TSG101 Cells | 121 |

B)

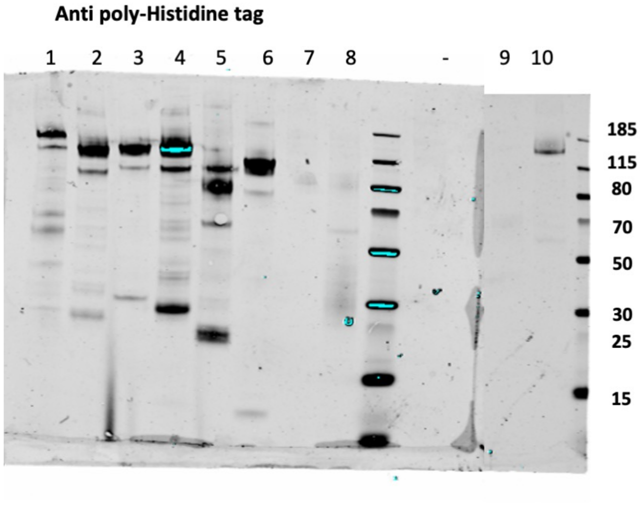


| Lane | Sample Name | Predicted M.W kDa |
|---|---|---|
| 1 | IL6ST-3X N term Syntenin EVs | 118 |
| 2 | IL6ST-N term Syntenin EVs | 91 |
| 3 | IL6ST-Syndecan1 EVs | 93 |
| 4 | IL6ST-GCN4 L.Z-N term Syntenin EVs | 94 |
| 5 | IL6ST extracellular domain-PDGFR TM-N term Syntenin EVs | 57 |
| 6 | IL6ST EVs | 77 |
| 7 | IL6ST extracellular domain-ADRB2 TM-CD63 EVs | 70 |
| 8 | Ctrl EVs | |
| 9 | IL6ST extracellular domain-PDGFR TM EVs | 43 |
| 10 | IL6ST-TSG101 EVs | 121 |

**Supplementary Figure 5.** WB probed against poly Histidine tag on **A)** engineered HEK293T cells and their **B)** respective EVs. Cells were transfected with indicated genetic constructs. Predicted molecular weight (kDa) values were calculated based on the amino acid sequence of the respective construct.

#### Supplementary Figure 6

A)

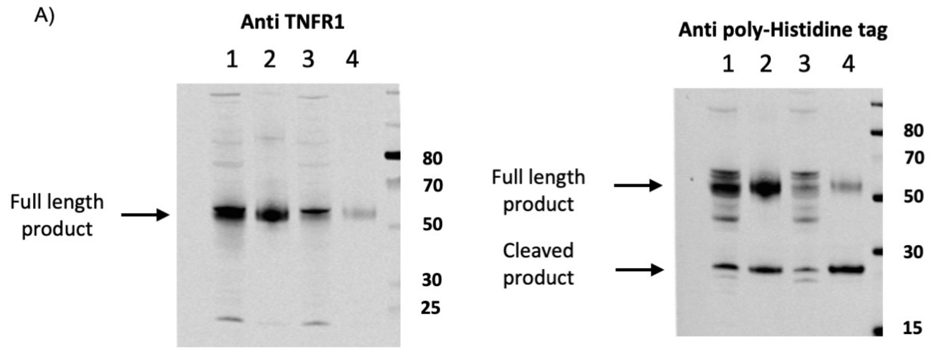


| Lane | Sample Name | Predicted M.W kDa |
|---|---|---|
| 1 | TNFR1 MMP-N term Syntenin Cells | 45 |
| 2 | TNFR1 MMP-N term Syntenin EVs | 45 |
| 3 | TNFR1-N term Syntenin Cells | 45 |
| 4 | TNFR1-N term Syntenin EVs | 45 |

B)

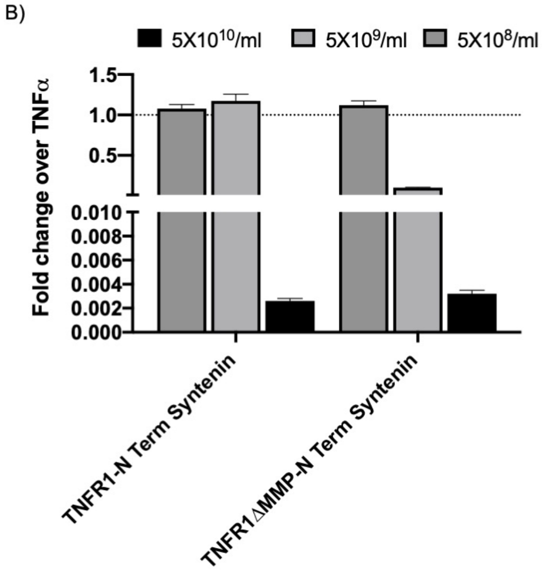


**Supplementary Figure 6. A)** WB probed against human TNFR1 and poly Histidine tag on engineered HEK293T EVs and their respective cells transfected with WT and Mutant TNFR1 fused to Syntenin. Predicted molecular weight (kDa) values were calculated based on the amino acid sequence of the respective construct. **B)** Engineered EVs displaying TNFR1 purified from HEK293T cells transfected with constructs listed in the figure legends, evaluated for TNFα decoy in an *in vitro* cell assay respondent to TNFα induced NF-ϰB activation. Data were normalized to control cells treated with TNFα (5ng/ml). **** $P < 0.0001$ statistical significance calculated by two-way ANOVA upon comparing response to Ctrl EVs at the respective dose.

#### Supplementary Figure 7

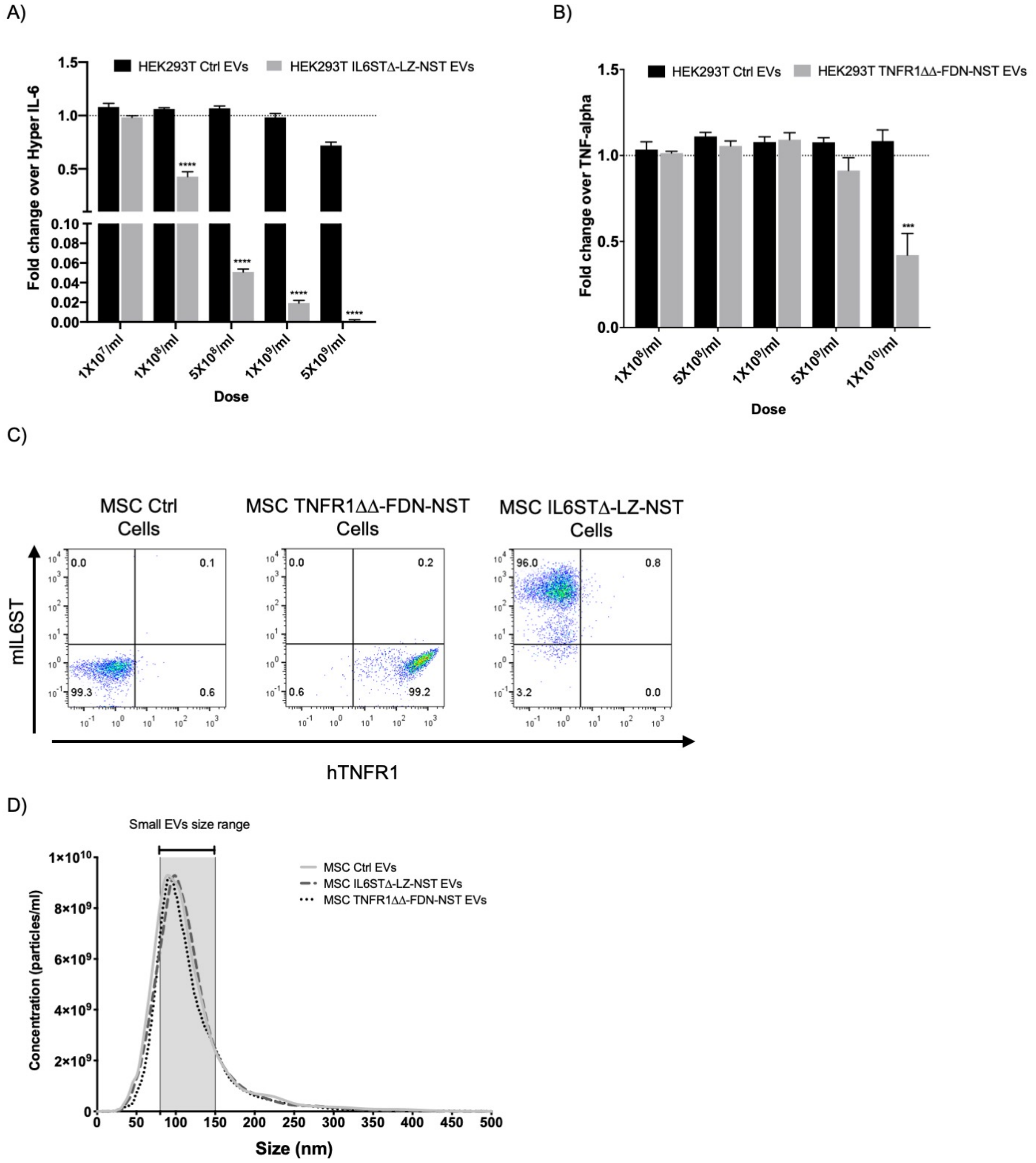


**Supplementary Figure 7. A)** IL6/IL6R induced STAT3 activation is reduced in a dose dependent manner by EVs purified from HEK293T cells stable expressing the optimized IL6STΔ-LZ-NST display construct. Data were normalized to control cells treated with IL6/sIL6R (5 ng/ml). Error bars, s.d. (*n*= 3). **** *P* < 0.0001 statistical significance calculated by two-way ANOVA upon comparing response to Ctrl EVs at the respective dose. **B)** TNFα induced NF-κB activation is reduced in a dose dependent manner by EVs purified from HEK293T cells stable expressing the optimized TNFR1ΔΔ-FDN-NST display construct. Data were normalized to control cells treated with TNFα (5ng/ml). Error bars, s.d. (*n*=3). **** PI < 0.0001 statistical significance calculated by two-way ANOVA upon comparing response to Ctrl EVs at the respective dose. **C)** Flow cytometry analysis of MSC TNFR1ΔΔ-FDN-NST, MSC IL6STΔ-LZ-NST cells stained with mouse IL6ST APC conjugated (y axis) and human TNFR1 PE (y axis) conjugated antibody. D) Size distribution determined by NTA of MSC Ctrl EVs, IL6STΔ-LZ-NST EVs and TNFR1ΔΔ-FDN-NST EVs.

#### Supplementary Figure 8

A)

**Anti poly-Histidine tag**

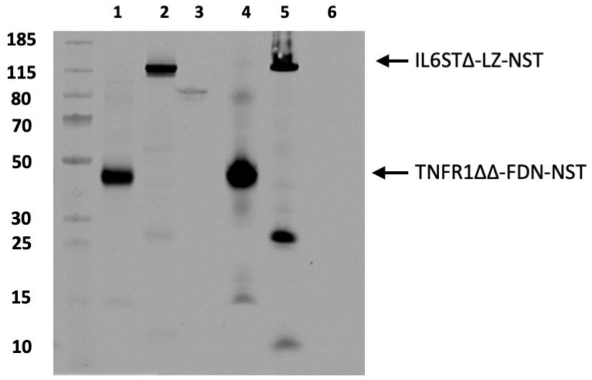


| Lane | Sample Name | Predicted M.W kDa |
|---|---|---|
| 1 | MSC TNFR1ΔΔ-FDN-NST Cells | 48 |
| 2 | MSC IL6STΔ-LZ-NST Cells | 94 |
| 3 | MSC Ctrl Cells | - |
| 4 | MSC TNFR1ΔΔ-FDN-NST EVs | 48 |
| 5 | MSC IL6STΔ-LZ-NST EVs | 94 |
| 6 | MSC Ctrl EVs | - |

B)

**Anti ALIX (Predicted MW kDa: 96)**

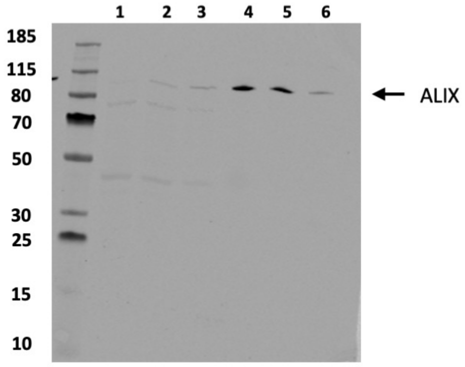


| Lane | Sample Name |
|---|---|
| 1 | MSC TNFR1ΔΔ-FDN-NST Cells |
| 2 | MSC IL6STΔ-LZ-NST Cells |
| 3 | MSC Ctrl Cells |
| 4 | MSC TNFR1ΔΔ-FDN-NST EVs |
| 5 | MSC IL6STΔ-LZ-NST EVs |
| 6 | MSC Ctrl EVs |

C)

**Anti Calnexin (Predicted MW kDa: 67) and Anti TSG101 (Predicted MW kDa: 44)**

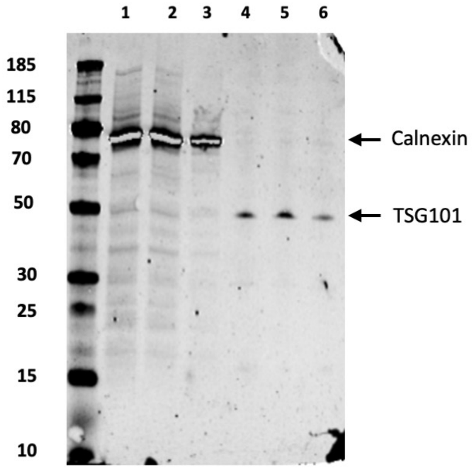


| Lane | Sample Name |
|---|---|
| 1 | MSC TNFR1ΔΔ-FDN-NST Cells |
| 2 | MSC IL6STΔ-LZ-NST Cells |
| 3 | MSC Ctrl Cells |
| 4 | MSC TNFR1ΔΔ-FDN-NST EVs |
| 5 | MSC IL6STΔ-LZ-NST EVs |
| 6 | MSC Ctrl EVs |

**Supplementary Figure 8. A-C)** Uncropped version of WB images as shown in Figure 3D.

### Supplementary Figure 9

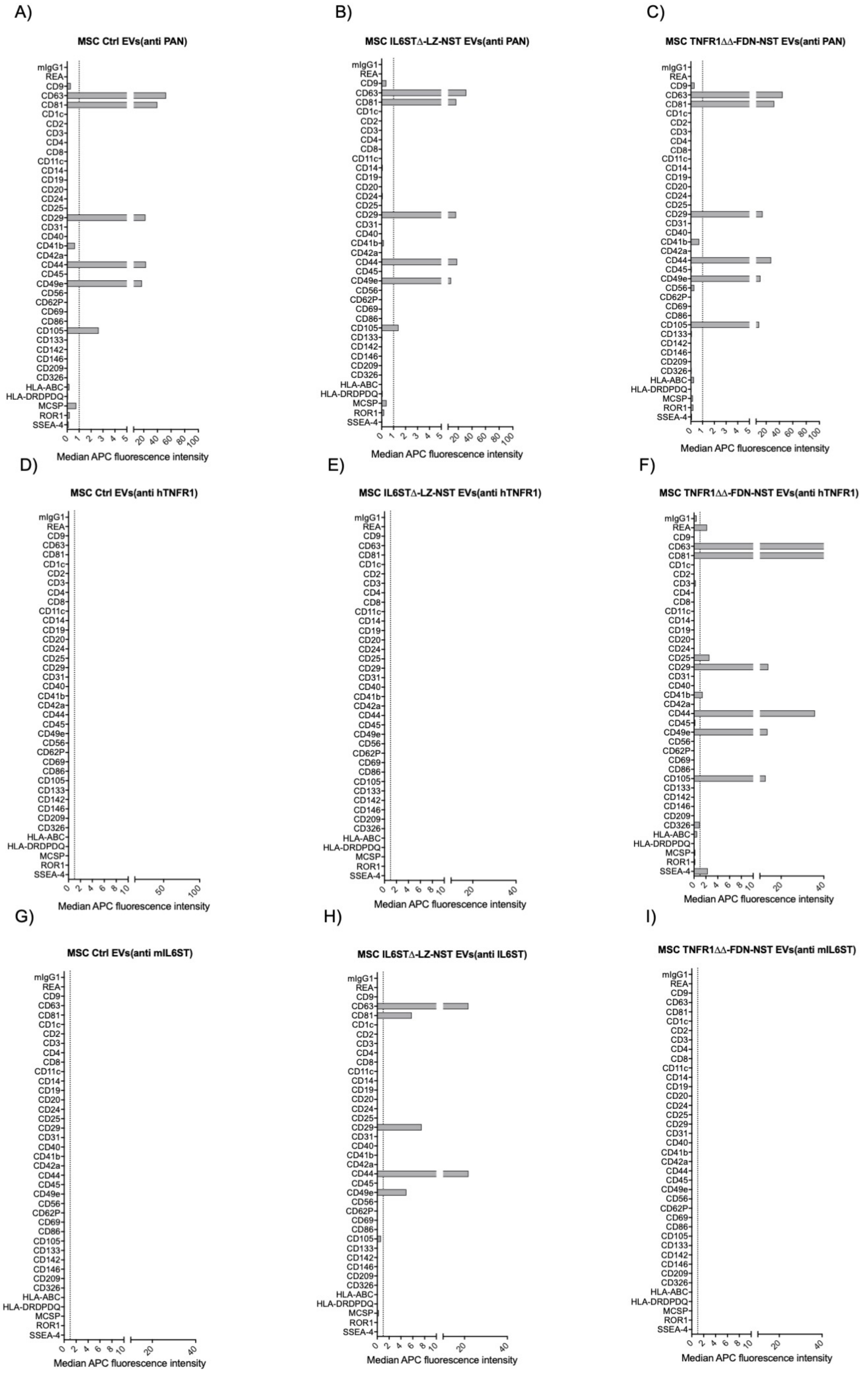

**Supplementary Figure 9. A-I)** Full expression profile of all the 37 different markers determined by multiplex bead-based assay for the same dataset as shown in Figure 3F-H.

#### Supplementary Figure 10

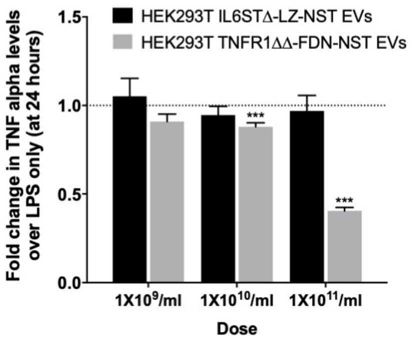


**Supplementary Figure 10.** Effect of HEK293T TNFR1ΔΔ-FDN-NST EVs and IL6STΔ-LZ-NST EVs on TNFα levels in conditioned medium determined by ELISA at 24 hours post LPS stimulation of RAW 246.7 macrophages. Data were normalized to control cells treated with LPS only. Error bars, SEM ($n$=3). *** $P < 0.0001$, statistical significance calculated by one-way ANOVA upon comparing response to Ctrl EVs at the respective dose.

#### Supplementary Figure 11

A)

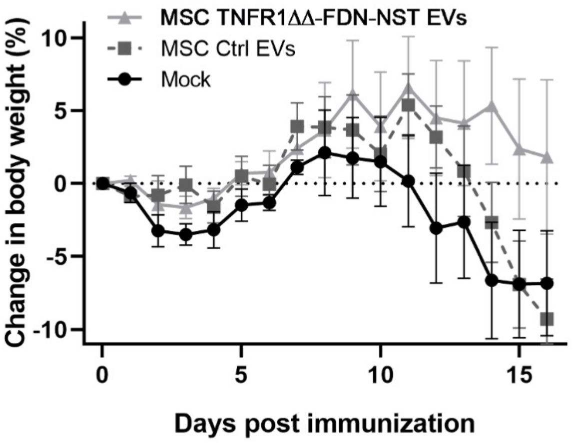


B)

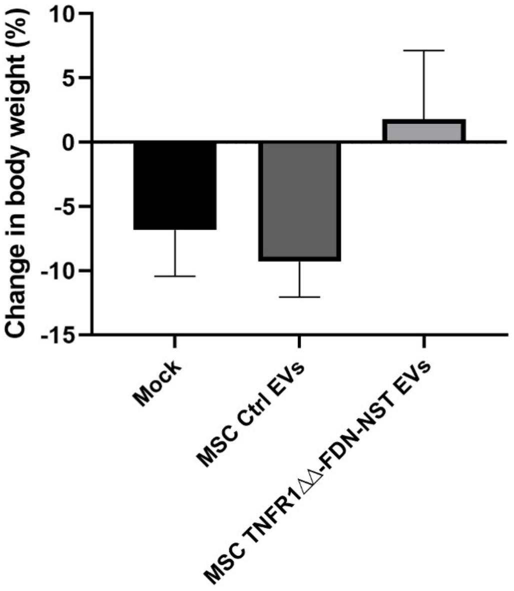


**Supplementary Figure 11. A)** Percent change in relative bodyweight to initial weight over the disease course and **B)** Percent change in body weight between day 0 to day 16 in mice induced with EAE using $MOG_{35-55}$ peptide and treated with S.C administration of either $4\times10^{10}$ MSC TNFR1ΔΔ-FDN-NST EVs (*n*=5) or MSC Ctrl EVs (*n*=5) or Saline (*n*=5) (On day 7, 10 & 13).

Supplementary Figure 12

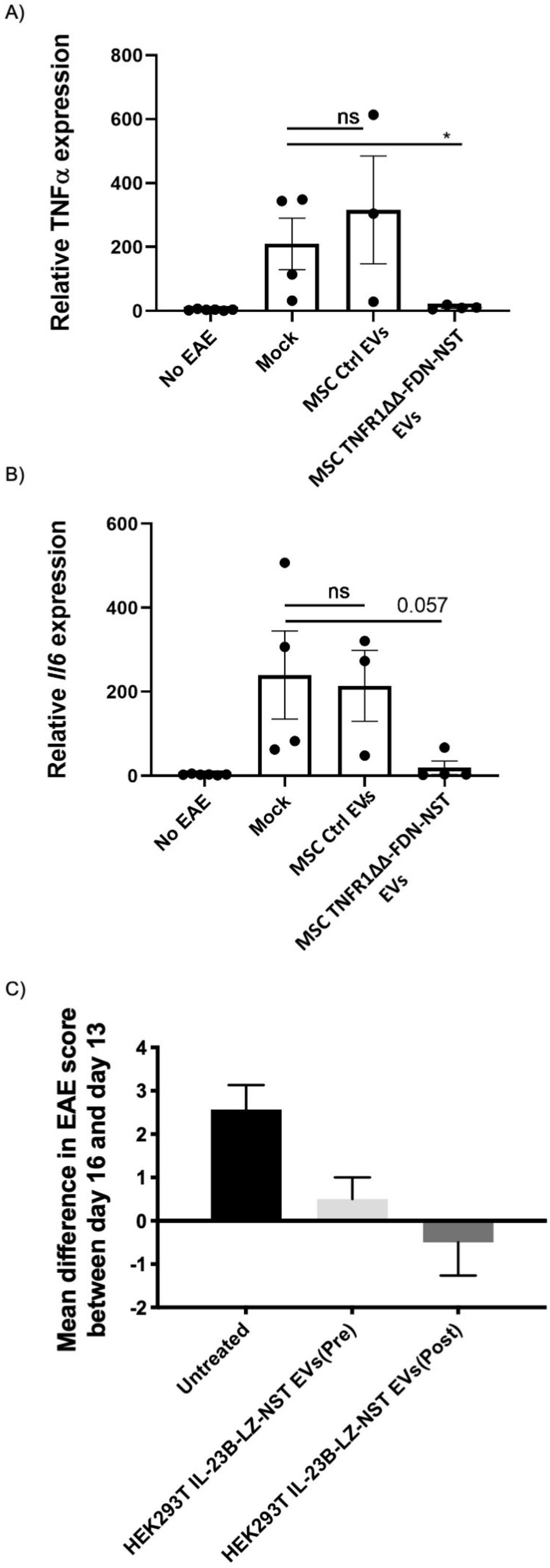


**Supplementary Figure 12.** Relative mRNA expression of pro inflammatory cytokines determined by qPCR **(A)** TNFα, **(B)** IL6 in the spinal cord at day 16 in mice induced with EAE using $MOG_{35-55}$ peptide and treated with S.C administration of either $4{\times}10^{10}$ MSC TNFR1ΔΔ-FDN-NST EVs (*n*=5) or MSC Ctrl EVs (*n*=5) or Saline (*n*=5) (On day 7, 10 & 13). For the same dataset as shown in Figure 5B-C**.** **C)** Relative change in clinical score of disease progression between day 16 and 13 in mice induced with EAE using $MOG_{35-55}$ peptide and treated with I.V administration of either $1{\times}10^{10}$ HEK293T IL23B-LZ-NST EVs pre symptomatic (On day 5, 7 & 10) or $6{\times}10^{10}$ HEK293T IL23B-LZ-NST EVs post symptomatic (On day 13) or Saline. For the same dataset as shown in Figure 5H-I.

#### Supplementary Figure 13

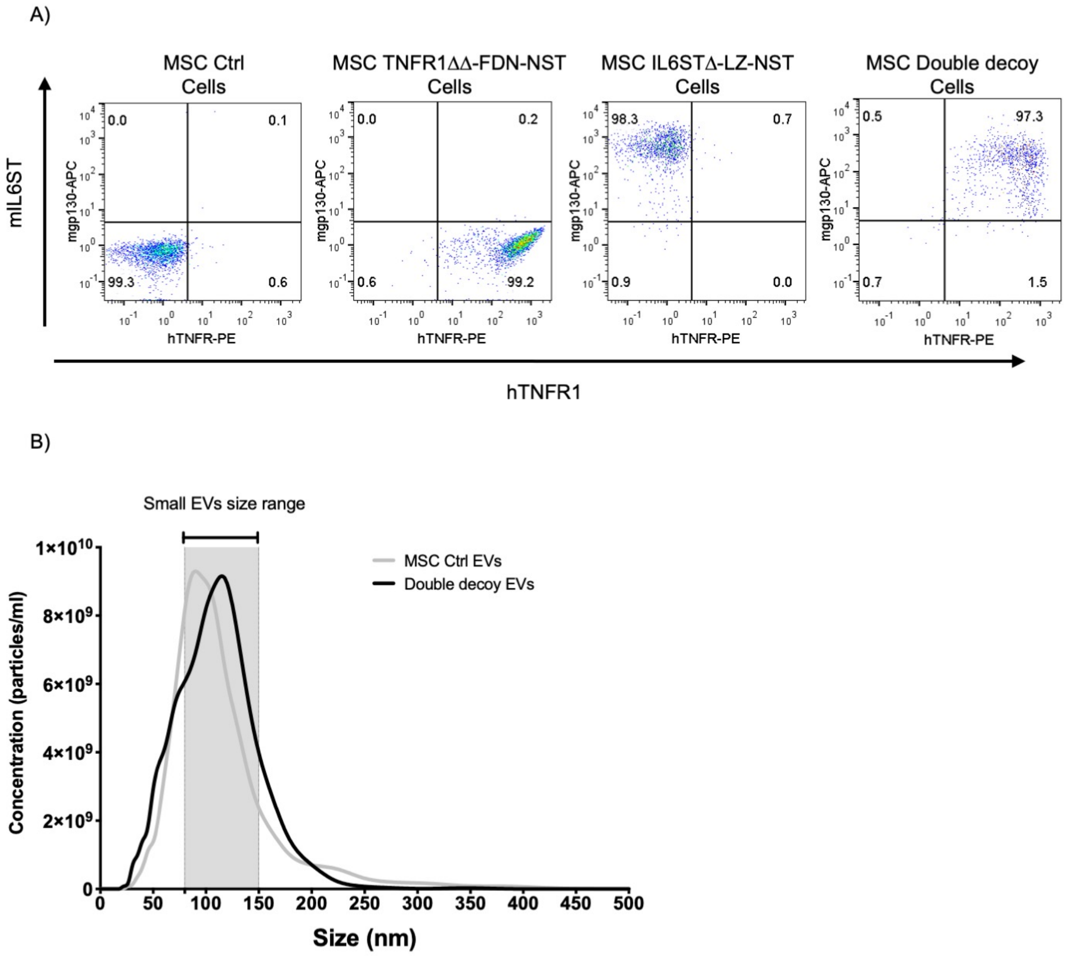


**Supplementary Figure 13. A)** Flow cytometry analysis of MSC TNFR1ΔΔ-FDN-NST, MSC IL6STΔ-LZ-NST and MSC double decoy cells stained with mouse IL6ST APC conjugated and human TNFR1 PE conjugated antibody. **B)** Size distribution determined by NTA of MSC Ctrl EVs and double decoy EVs.

#### Supplementary Figure 14

A)

**Anti poly-Histidine tag**

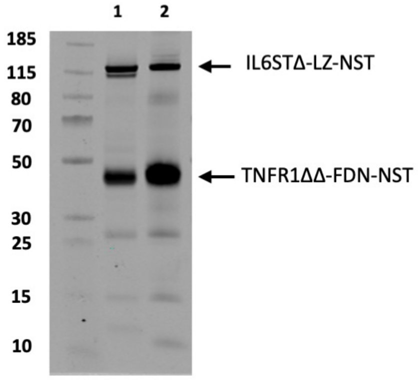


| Lane | Sample Name | Predicted M.W kDa |
|---|---|---|
| 1 | MSC Double decoy Cells | 94 and 48 |
| 2 | MSC Double decoy EVs | 94 and 48 |

B)

**Anti ALIX (Predicted MW kDa: 96)**

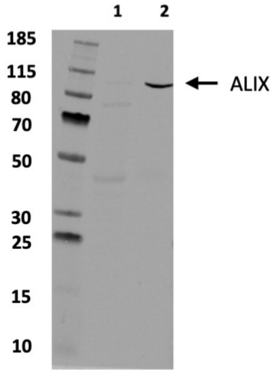


| Lane | Sample Name |
|---|---|
| 1 | MSC Double decoy Cells |
| 2 | MSC Double decoy EVs |

C)

**Anti Calnexin (Predicted MW kDa: 67) and Anti TSG101 (Predicted MW kDa: 44)**

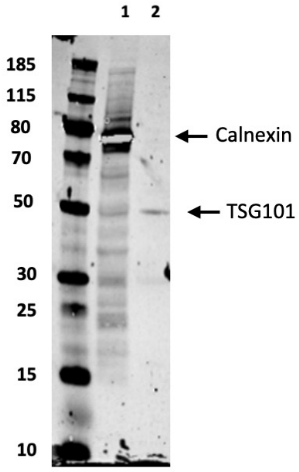


| Lane | Sample Name |
|---|---|
| 1 | MSC Double decoy Cells |
| 2 | MSC Double decoy EVs |

**Supplementary Figure 14. A)** WB of double decoy cells and EVs indicating the presence of both his-tagged decoy receptors; TNFR1ΔΔ-FDN-NST (48 kDa) and IL6STΔ-LZ-NST (94 kDa) in cells and EVs. The WB results further demonstrate the presence of **B)** classical EV markers; ALIX (96 kDa), TSG101 (44 kDa) and **C)** absence of Calnexin (67 kDa) in the isolated EVs.

#### Supplementary Figure 15

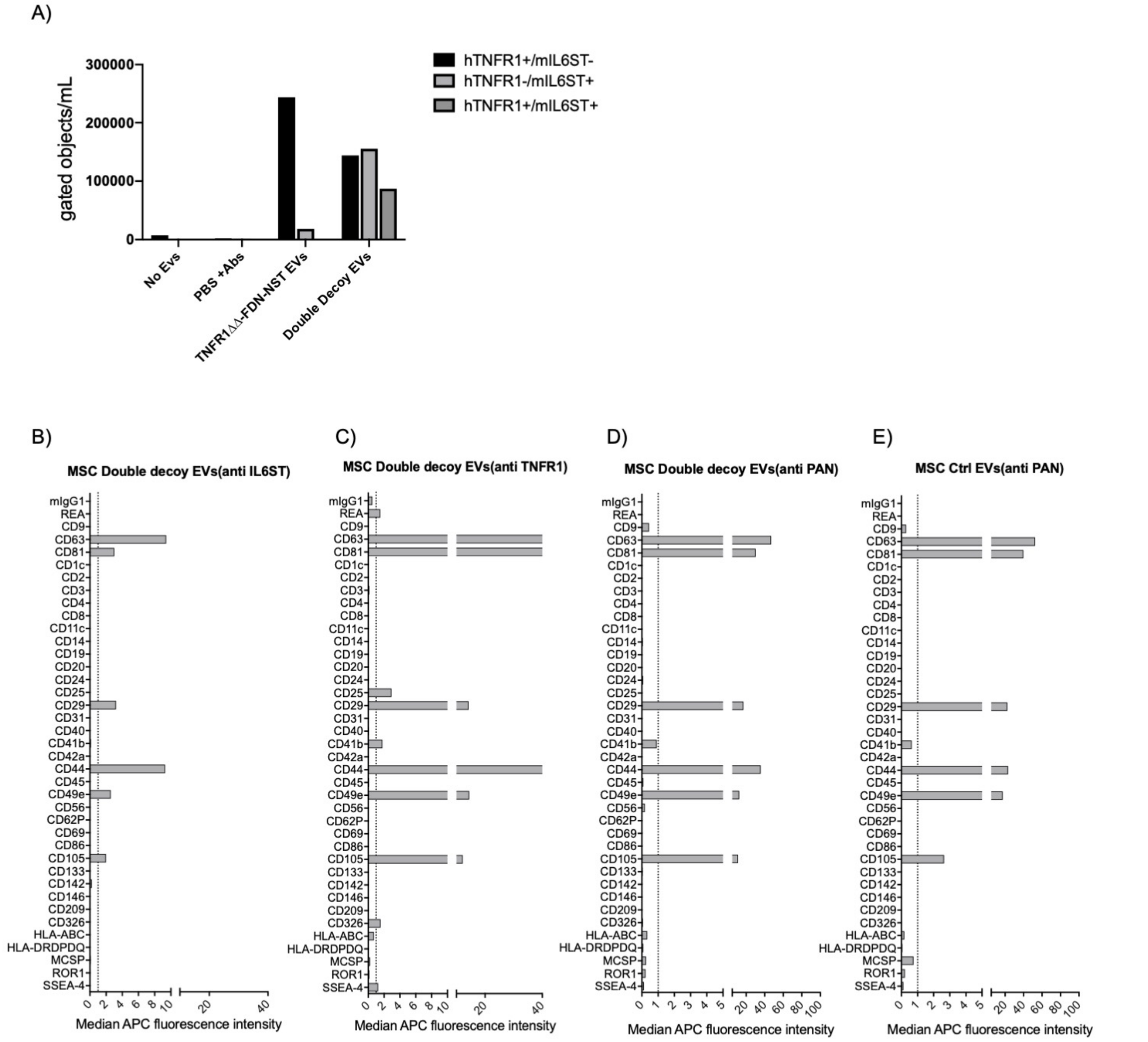


**Supplementary Figure 15 A)** Concentration of events in different gated population, for the same dataset as shown in Figure 6A-B. **B-E).** Full expression profile of all the 37 different markers determined by multiplex bead-based assay for the same dataset as shown in Figure 6D.

#### Supplementary Figure 16

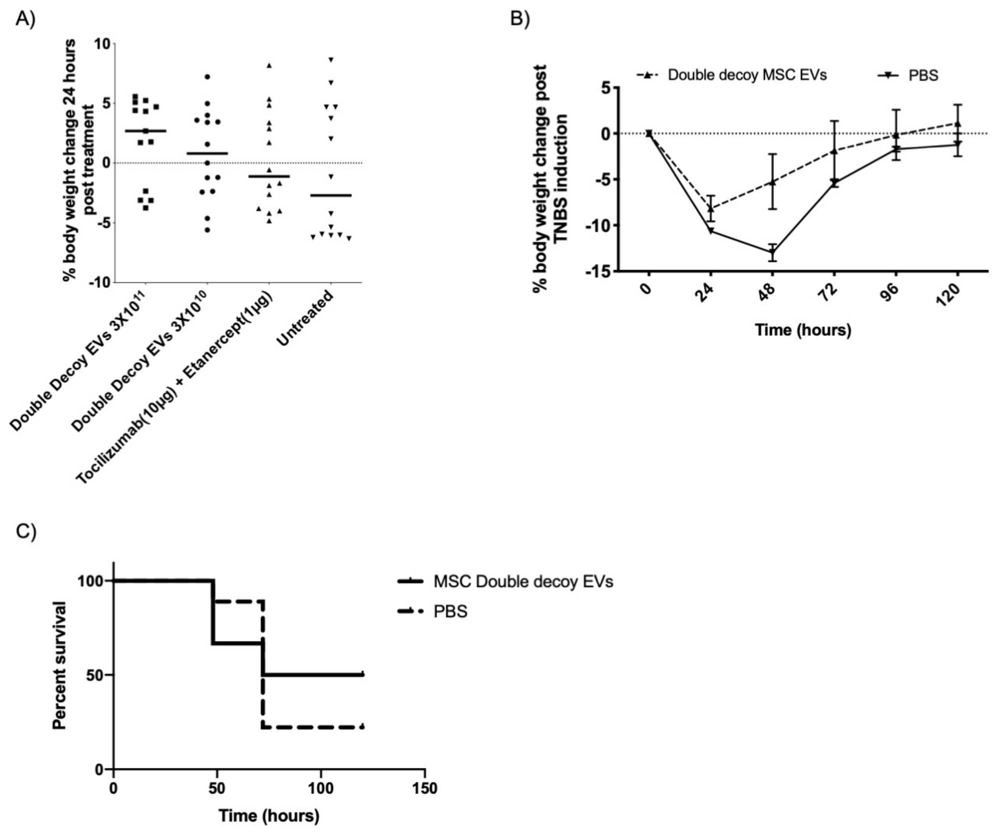


**Supplementary Figure 16. A)** Change in body weight 24 hours after treatment with different groups in mice induced with TNBS colitis. For the same data set as shown in Figure 6G-H. **B)** Survival curve and **C)** percent change in relative bodyweight to initial weight over the disease course in mice induced with colitis by intrarectal injection of TNBS and treated with I.V administration (at 24 hours post disease induction) of either $3\times10^{11}$ MSC double decoy EVs (*n*=8) or PBS (*n*=9).

**Supplementary Table 1**

| S.No | Construct name | Amino acid sequence |
|---|---|---|
| 1 | **mIL6ST TSG101** | **MSAPRIWLAQALLFFLTTESIGQLLEPCGYIYPEFPVVQRGSNFTAICVLKEACLQHYYVN<br>ASYIVWKTNHAAVPREQVTVINRTTSSVTFTDVVLPSVQLTCNILSFGQIEQNVYGVTML<br>SGFPPDKPTNLTCIVNEGKNMLCQWDPGRETYLETNYTLKSEWATEKFPDCQSKHGTS<br>CMVSYMPTYYVNIEVWVEAENALGKVSSESINFDPVDKVKPTPPYNLSVTNSEELSSILK<br>LSWVSSGLGGLLDLKSDIQYRTKDASTWIQVPLEDTMSPRTSFTVQDLKPFTEYVFRIRSI<br>KDSGKGYWSDWSEEASGTTYEDRPSRPPSFWYKTNPSHGQEYRSVRLIWKALPLSEAN<br>GKILDYEVILTQSKSVSQTYTVTGTELTVNLTNDRYVASLAARNKVGKSAAAVLTIPSPHV<br>TAAYSVVNLKAFPKDNLLWVEWTPPPKPVSKYILEWCVLSENAPCVEDWQQEDATVN<br>RTHLRGRLLESKCYQITVTPVFATGPGGSESLKAYLKQAAPARGPTVRTKKVGKNEAVLA<br>WDQIPVDDQNGFIRNYSISYRTSVGKEMVVHVDSSHTEYTLSSLSSDTLYMVRMAAYT<br>DEGGKDGPEFTFTTPKFAQGEIEAIVVPVCLAFLLTTLLGVLFCFNKRDLIKKHIWPNVPD<br>PSKSHIAQWSPHTPPRHNFNSKDQGSGSGSGSGSAVSESQLKKMVSKYKYRDLTVRET<br>VNVITLYKDLKPVLDSYVFNDGSSRELMNLTGTIPVPYRGNTYNIPICLWLLDTYPYNPPI<br>CFVKPTSSMTIKTGKHVDANGKIYLPYLHEWKHPQSDLLGLIQVMIVVFGDEPPVFSRPI<br>SASYPPYQATGPPNTSYMPGMPGGISPYPSGYPPNPSGYPGCPYPPGGPYPATTSSQY<br>PSQPPVTTVGPSRDGTISEDTIRASLISAVSDKLRWRMKEEMDRAQAELNALKRTEEDL<br>KKGHQKLEEMVTRLDQEVAEVDKNIELLKKKDEELSSALEKMENQSENNDIDEVIIPTAP<br>LYKQILNLYAEENAIEDTIFYLGEALRRGVIDLDVFLKHVRLLSRKQFQLRALMQKARKTA<br>GLSDLYHHHHHH** |
| 2 | **mIL6ST N-term-Syntenin** | **MSAPRIWLAQALLFFLTTESIGQLLEPCGYIYPEFPVVQRGSNFTAICVLKEACLQHYYVN<br>ASYIVWKTNHAAVPREQVTVINRTTSSVTFTDVVLPSVQLTCNILSFGQIEQNVYGVTML<br>SGFPPDKPTNLTCIVNEGKNMLCQWDPGRETYLETNYTLKSEWATEKFPDCQSKHGTS<br>CMVSYMPTYYVNIEVWVEAENALGKVSSESINFDPVDKVKPTPPYNLSVTNSEELSSILK<br>LSWVSSGLGGLLDLKSDIQYRTKDASTWIQVPLEDTMSPRTSFTVQDLKPFTEYVFRIRSI<br>KDSGKGYWSDWSEEASGTTYEDRPSRPPSFWYKTNPSHGQEYRSVRLIWKALPLSEAN<br>GKILDYEVILTQSKSVSQTYTVTGTELTVNLTNDRYVASLAARNKVGKSAAAVLTIPSPHV<br>TAAYSVVNLKAFPKDNLLWVEWTPPPKPVSKYILEWCVLSENAPCVEDWQQEDATVN<br>RTHLRGRLLESKCYQITVTPVFATGPGGSESLKAYLKQAAPARGPTVRTKKVGKNEAVLA<br>WDQIPVDDQNGFIRNYSISYRTSVGKEMVVHVDSSHTEYTLSSLSSDTLYMVRMAAYT<br>DEGGKDGPEFTFTTPKFAQGEIEAIVVPVCLAFLLTTLLGVLFCFNKRDLIKKHIWPNVPD<br>PSKSHIAQWSPHTPPRHNFNSKDQGGGGSGGGGSGGGGSSLYPSLEDLKVDKVIQAQ<br>TAYSANPASQAFVLVDASAALPPDGNLYPKLYPELSQYMGLSLNEAEICESMPMVSGAP<br>AQGQLVARPSSVNYMVAPVTGNDAGIRRAEIKHHHHHH** |

| | | |
|---|---|---|
| 3 | **mIL6ST** | **MSAPRIWLAQALLFFLTTESIGQLLEPCGYIYPEFPVVQRGSNFTAICVLKEACLQHYYVN ASYIVWKTNHAAVPREQVTVINRTTSSVTFTDVVLPSVQLTCNILSFGQIEQNVYGVTML SGFPPDKPTNLTCIVNEGKNMLCQWDPGRETYLETNYTLKSEWATEKFPDCQSKHGTS CMVSYMPTYYVNIEVWVEAENALGKVSSESINFDPVDKVKPTPPYNLSVTNSEELSSILK LSWVSSGLGGLLDLKSDIQYRTKDASTWIQVPLEDTMSPRTSFTVQDLKPFTEYVFRIRSI KDSGKGYWSDWSEEASGTTYEDRPSRPPSFWYKTNPSHGQEYRSVRLIWKALPLSEAN GKILDYEVILTQSKSVSQTYTVTGTELTVNLTNDRYVASLAARNKVGKSAAAVLTIPSPHV TAAYSVVNLKAFPKDNLLWVEWTPPPKPVSKYILEWCVLSENAPCVEDWQQEDATVN RTHLRGRLLESKCYQITVTPVFATGPGGSESLKAYLKQAAPARGPTVRTKKVGKNEAVLA WDQIPVDDQNGFIRNYSISYRTSVGKEMVVHVDSSHTEYTLSSLSSDTLYMVRMAAYT DEGGKDGPEFTFTTPKFAQGEIEAIVVPVCLAFLLTTLLGVLFCFNKRDLIKKHIWPNVPD PSKSHIAQWSPHTPPRHNFNSKDQHHHHHH** |
| 4 | **mIL6ST Syndecan 1 x 3** | **MSAPRIWLAQALLFFLTTESIGQLLEPCGYIYPEFPVVQRGSNFTAICVLKEACLQHYYVN ASYIVWKTNHAAVPREQVTVINRTTSSVTFTDVVLPSVQLTCNILSFGQIEQNVYGVTML SGFPPDKPTNLTCIVNEGKNMLCQWDPGRETYLETNYTLKSEWATEKFPDCQSKHGTS CMVSYMPTYYVNIEVWVEAENALGKVSSESINFDPVDKVKPTPPYNLSVTNSEELSSILK LSWVSSGLGGLLDLKSDIQYRTKDASTWIQVPLEDTMSPRTSFTVQDLKPFTEYVFRIRSI KDSGKGYWSDWSEEASGTTYEDRPSRPPSFWYKTNPSHGQEYRSVRLIWKALPLSEAN GKILDYEVILTQSKSVSQTYTVTGTELTVNLTNDRYVASLAARNKVGKSAAAVLTIPSPHV TAAYSVVNLKAFPKDNLLWVEWTPPPKPVSKYILEWCVLSENAPCVEDWQQEDATVN RTHLRGRLLESKCYQITVTPVFATGPGGSESLKAYLKQAAPARGPTVRTKKVGKNEAVLA WDQIPVDDQNGFIRNYSISYRTSVGKEMVVHVDSSHTEYTLSSLSSDTLYMVRMAAYT DEGGKDGPEFTFTTPKFAQGEIEAIVVPVCLAFLLTTLLGVLFCFNKRDLIKKHIWPNVPD PSKSHIAQWSPHTPPRHNFNSKDQGGGGSGGGGSGGGGSRMKKKDEGSYSLEEPKQ ANGGAYQKPTKQEEFYAGGGGSGGGGSGGGGSRMKKKDEGSYSLEEPKQANGGAYQ KPTKQEEFYAGGGGSGGGGSGGGGSRMKKKDEGSYSLEEPKQANGGAYQKPTKQEEF YAHHHHHH** |
| 5 | **mIL6ST-GCN4 LZ-N-term-Syntenin** | **MSAPRIWLAQALLFFLTTESIGQLLEPCGYIYPEFPVVQRGSNFTAICVLKEACLQHYYVN ASYIVWKTNHAAVPREQVTVINRTTSSVTFTDVVLPSVQLTCNILSFGQIEQNVYGVTML SGFPPDKPTNLTCIVNEGKNMLCQWDPGRETYLETNYTLKSEWATEKFPDCQSKHGTS CMVSYMPTYYVNIEVWVEAENALGKVSSESINFDPVDKVKPTPPYNLSVTNSEELSSILK LSWVSSGLGGLLDLKSDIQYRTKDASTWIQVPLEDTMSPRTSFTVQDLKPFTEYVFRIRSI KDSGKGYWSDWSEEASGTTYEDRPSRPPSFWYKTNPSHGQEYRSVRLIWKALPLSEAN GKILDYEVILTQSKSVSQTYTVTGTELTVNLTNDRYVASLAARNKVGKSAAAVLTIPSPHV TAAYSVVNLKAFPKDNLLWVEWTPPPKPVSKYILEWCVLSENAPCVEDWQQEDATVN RTHLRGRLLESKCYQITVTPVFATGPGGSESLKAYLKQAAPARGPTVRTKKVGKNEAVLA WDQIPVDDQNGFIRNYSISYRTSVGKEMVVHVDSSHTEYTLSSLSSDTLYMVRMAAYT DEGGKDGPEFTFTTPKFAQGEIEAIVVPVCLAFLLTTLLGVLFCFNKRDLIKKHIWPNVPD PSKSHIAQWSPHTPPRHNFNSKDQRMKQLEDKVEELLSKNYHLENEVARLKKLVGERG SGSGSGSGSSLYPSLEDLKVDKVIQAQTAYSANPASQAFVLVDASAALPPDGNLYPKLYP ELSQYMGLSLNEAEICESMPMVSGAPAQGQLVARPSSVNYMVAPVTGNDAGIRRAEI KHHHHHH** |

| | | |
|---|---|---|
| 6 | **mIL6ST PDGFR TM N-term-Syntenin** | **MSAPRIWLAQALLFFLTTESIGQLLEPCGYIYPEFPVVQRGSNFTAICVLKEACLQHYYVN<br>ASYIVWKTNHAAVPREQVTVINRTTSSVTFTDVVLPSVQLTCNILSFGQIEQNVYGVTML<br>SGFPPDKPTNLTCIVNEGKNMLCQWDPGRETYLETNYTLKSEWATEKFPDCQSKHGTS<br>CMVSYMPTYYVNIEVWVEAENALGKVSSESINFDPVDKVKPTPPYNLSVTNSEELSSILK<br>LSWVSSGLGGLLDLKSDIQYRTKDASTWIQVPLEDTMSPRTSFTVQDLKPFTEYVFRIRSI<br>KDSGKGYWSDWSEEASGTTYEDSTASFAVGQDTQEVIVVPHSLPFKVVVISAILALVVLT<br>IISLIILIMLWQKKPRGSGSGSGSGSSLYPSLEDLKVDKVIQAQTAYSANPASQAFVLVDA<br>SAALPPDGNLYPKLYPELSQYMGLSLNEAEICESMPMVSGAPAQGQLVARPSSVNYMV<br>APVTGNDAGIRRAEIKHHHHHHHHHHHH** |
| 7 | **mIL6ST ADRB2 TM CD63** | **MSAPRIWLAQALLFFLTTESIGQLLEPCGYIYPEFPVVQRGSNFTAICVLKEACLQHYYVN<br>ASYIVWKTNHAAVPREQVTVINRTTSSVTFTDVVLPSVQLTCNILSFGQIEQNVYGVTML<br>SGFPPDKPTNLTCIVNEGKNMLCQWDPGRETYLETNYTLKSEWATEKFPDCQSKHGTS<br>CMVSYMPTYYVNIEVWVEAENALGKVSSESINFDPVDKVKPTPPYNLSVTNSEELSSILK<br>LSWVSSGLGGLLDLKSDIQYRTKDASTWIQVPLEDTMSPRTSFTVQDLKPFTEYVFRIRSI<br>KDSGKGYWSDWSEEASGTTYEDRPSRPPSFWYKTNPSHGQEYRSVRLIWKALPLSEAN<br>GKILDYEVILTQSKSVSQTYTVTGTELTVNLTNDRYVASLAARNKVGKSAAAVLTIPSPHV<br>TAAYSVVNLKAFPKDNLLWVEWTPPPKPVSKYILEWCVLSENAPCVEDWQQEDATVN<br>RTHLRGRLLESKCYQITVTPVFATGPGGSESLKAYLKQAAPARGPTVRTKKVGKNEAVLA<br>WDQIPVDDQNGFIRNYSISYRTSVGKEMVVHVDSSHTEYTLSSLSSDTLYMVRMAAYT<br>DEGGKDGPEFTFTTPKFASESPEPLSQQWTAGMGLLMALIVLLIVAGNVLVIVAIAKTPR<br>LAVEGGMKCVKFLLYVLLLAFCACAVGLIAVGVGAQLVLSQTIIQGATPGSLLPVVIIAVG<br>VFLFLVAFVGCCGACKENYCLMITFAIFLSLIMLVEVAAAIAGYVFRDKVMSEFNNNFRQ<br>QMENYPKNNHTASILDRMQADFKCCGAANYTDWEKIPSMSKNRVPDSCCINVTVGC<br>GINFNEKAIHKEGCVEKIGGWLRKNVLVVAAAALGIAFVEVLGIVFACCLVKSIRSGYEV<br>MHHHHHH*** |
| 8 | **mIL6ST N-term-Syntenin X 3** | **MSAPRIWLAQALLFFLTTESIGQLLEPCGYIYPEFPVVQRGSNFTAICVLKEACLQHYYVN<br>ASYIVWKTNHAAVPREQVTVINRTTSSVTFTDVVLPSVQLTCNILSFGQIEQNVYGVTML<br>SGFPPDKPTNLTCIVNEGKNMLCQWDPGRETYLETNYTLKSEWATEKFPDCQSKHGTS<br>CMVSYMPTYYVNIEVWVEAENALGKVSSESINFDPVDKVKPTPPYNLSVTNSEELSSILK<br>LSWVSSGLGGLLDLKSDIQYRTKDASTWIQVPLEDTMSPRTSFTVQDLKPFTEYVFRIRSI<br>KDSGKGYWSDWSEEASGTTYEDRPSRPPSFWYKTNPSHGQEYRSVRLIWKALPLSEAN<br>GKILDYEVILTQSKSVSQTYTVTGTELTVNLTNDRYVASLAARNKVGKSAAAVLTIPSPHV<br>TAAYSVVNLKAFPKDNLLWVEWTPPPKPVSKYILEWCVLSENAPCVEDWQQEDATVN<br>RTHLRGRLLESKCYQITVTPVFATGPGGSESLKAYLKQAAPARGPTVRTKKVGKNEAVLA<br>WDQIPVDDQNGFIRNYSISYRTSVGKEMVVHVDSSHTEYTLSSLSSDTLYMVRMAAYT<br>DEGGKDGPEFTFTTPKFAQGEIEAIVVPVCLAFLLTTLLGVLFCFNKRDLIKKHIWPNVPD<br>PSKSHIAQWSPHTPPRHNFNSKDQGGGGSGGGGSGGGGSSLYPSLEDLKVDKVIQAQ<br>TAYSANPASQAFVLVDASAALPPDGNLYPKLYPELSQYMGLSLNEAEICESMPMVSGAP<br>AQGQLVARPSSVNYMVAPVTGNDAGIRRAEIKGGGGSGGGGSGGGGSSLYPSLEDLK<br>VDKVIQAQTAYSANPASQAFVLVDASAALPPDGNLYPKLYPELSQYMGLSLNEAEICES<br>MPMVSGAPAQGQLVARPSSVNYMVAPVTGNDAGIRRAEIKGGGGSGGGGSGGGGS<br>SLYPSLEDLKVDKVIQAQTAYSANPASQAFVLVDASAALPPDGNLYPKLYPELSQYMGLS<br>LNEAEICESMPMVSGAPAQGQLVARPSSVNYMVAPVTGNDAGIRRAEIKHHHHHH*** |

| | | |
|---|---|---|
| 9 | **mIL6ST Flottlin 1** | **MSAPRIWLAQALLFFLTTESIGQLLEPCGYIYPEFPVVQRGSNFTAICVLKEACLQHYYVN<br>ASYIVWKTNHAAVPREQVTVINRTTSSVTFTDVVLPSVQLTCNILSFGQIEQNVYGVTML<br>SGFPPDKPTNLTCIVNEGKNMLCQWDPGRETYLETNYTLKSEWATEKFPDCQSKHGTS<br>CMVSYMPTYYVNIEVWVEAENALGKVSSESINFDPVDKVKPTPPYNLSVTNSEELSSILK<br>LSWVSSGLGGLLDLKSDIQYRTKDASTWIQVPLEDTMSPRTSFTVQDLKPFTEYVFRIRSI<br>KDSGKGYWSDWSEEASGTTYEDRPSRPPSFWYKTNPSHGQEYRSVRLIWKALPLSEAN<br>GKILDYEVILTQSKSVSQTYTVTGTELTVNLTNDRYVASLAARNKVGKSAAAVLTIPSPHV<br>TAAYSVVNLKAFPKDNLLWVEWTPPPKPVSKYILEWCVLSENAPCVEDWQQEDATVN<br>RTHLRGRLLESKCYQITVTPVFATGPGGSESLKAYLKQAAPARGPTVRTKKVGKNEAVLA<br>WDQIPVDDQNGFIRNYSISYRTSVGKEMVVHVDSSHTEYTLSSLSSDTLYMVRMAAYT<br>DEGGKDGPEFTFTTPKFAQGEIEAIVVPVCLAFLLTTLLGVLFCFNKRDLIKKHIWPNVPD<br>PSKSHIAQWSPHTPPRHNFNSKDQGSGSGSGSGSFFTCGPNEAMVVSGFCRSPPVMV<br>AGGRVFVLPCIQQIQRISLNTLTLNVKSEKVYTRHGVPISVTGIAQVKIQGQNKEMLAAA<br>CQMFLGKTEAEIAHIALETLEGHQRAIMAHMTVEEIYKDRQKFSEQVFKVASSDLVNM<br>GISVVSYTLKDIHDDQDYLHSLGKARTAQVQKDARIGEAEAKRDAGIREAKAKQEKVSA<br>QYLSEIEMAKAQRDYELKKAAYDIEVNTRRAQADLAYQLQVAKTKQQIEEQRVQVQVV<br>ERAQQVAVQEQEIARREKELEARVRKPAEAERYKLERLAEAEKSQLIMQAEAEAASVR<br>MRGEAEAFAIGARARAEAEQMAKKAEAFQLYQEAAQLDMLLEKLPQVAEEISGPLTSA<br>NKITLVSSGSGTMGAAKVTGEVLDILTRLPESVERLTGVSISQVNHKPLRTAHHHHHH*** |
| 10 | **mIL6ST-Fragment X-N-term-Syntenin** | **MSAPRIWLAQALLFFLTTESIGQLLEPCGYIYPEFPVVQRGSNFTAICVLKEACLQHYYVN<br>ASYIVWKTNHAAVPREQVTVINRTTSSVTFTDVVLPSVQLTCNILSFGQIEQNVYGVTML<br>SGFPPDKPTNLTCIVNEGKNMLCQWDPGRETYLETNYTLKSEWATEKFPDCQSKHGTS<br>CMVSYMPTYYVNIEVWVEAENALGKVSSESINFDPVDKVKPTPPYNLSVTNSEELSSILK<br>LSWVSSGLGGLLDLKSDIQYRTKDASTWIQVPLEDTMSPRTSFTVQDLKPFTEYVFRIRSI<br>KDSGKGYWSDWSEEASGTTYEDRPSRPPSFWYKTNPSHGQEYRSVRLIWKALPLSEAN<br>GKILDYEVILTQSKSVSQTYTVTGTELTVNLTNDRYVASLAARNKVGKSAAAVLTIPSPHV<br>TAAYSVVNLKAFPKDNLLWVEWTPPPKPVSKYILEWCVLSENAPCVEDWQQEDATVN<br>RTHLRGRLLESKCYQITVTPVFATGPGGSESLKAYLKQAAPARGPTVRTKKVGKNEAVLA<br>WDQIPVDDQNGFIRNYSISYRTSVGKEMVVHVDSSHTEYTLSSLSSDTLYMVRMAAYT<br>DEGGKDGPEFTFTTPKFAQGEIEAIVVPVCLAFLLTTLLGVLFCFNKRDLIKKHIWPNVPD<br>PSKSHIAQWSPHTPPRHNFNSKDQETIETFDNNEEESSYSYEEINDQTNDNITARLDRID<br>EKLSEILGMLHTLVVASAGPTSARGSGSGSGSGSSLYPSLEDLKVDKVIQAQTAYSANPA<br>SQAFVLVDASAALPPDGNLYPKLYPELSQYMGLSLNEAEICESMPMVSGAPAQGQLVA<br>RPSSVNYMVAPVTGNDAGIRRAEIKHHHHHH*** |

| | | |
|---|---|---|
| 11 | **mIL6ST-2XGCN4 LZ-N-term-Syntenin** | **MSAPRIWLAQALLFFLTTESIGQLLEPCGYIYPEFPVVQRGSNFTAICVLKEACLQHYYVN<br>ASYIVWKTNHAAVPREQVTVINRTTSSVTFTDVVLPSVQLTCNILSFGQIEQNVYGVTML<br>SGFPPDKPTNLTCIVNEGKNMLCQWDPGRETYLETNYTLKSEWATEKFPDCQSKHGTS<br>CMVSYMPTYYVNIEVWVEAENALGKVSSESINFDPVDKVKPTPPYNLSVTNSEELSSILK<br>LSWVSSGLGGLLDLKSDIQYRTKDASTWIQVPLEDTMSPRTSFTVQDLKPFTEYVFRIRSI<br>KDSGKGYWSDWSEEASGTTYEDRPSRPPSFWYKTNPSHGQEYRSVRLIWKALPLSEAN<br>GKILDYEVILTQSKSVSQTYTVTGTELTVNLTNDRYVASLAARNKVGKSAAAVLTIPSPHV<br>TAGSGSGSGSGSRMKQLEDKVEELLSKNYHLENEVARLKKLVGERGSGSGSGSGSDNKF<br>NKEQQNAFYEILHLPNLNEEQRNAFIQSLKDDPSQSANLLAEAKKLNDAQAPKAAPAR<br>GPTVRTKKVGKNEAVLAWDQIPVDDQNGFIRNYSISYRTSVGKEMVVHVDSSHTEYTL<br>SSLSSDTLYMVRMAAYTDEGGKDGPEFTFTTPKFAQGEIEAIVVPVCLAFLLTTLLGVLFC<br>FNKRDLIKKHIWPNVPDPSKSHIAQWSPHTPPRHNFNSKDQGSGSGSGSGSRMKQLE<br>DKVEELLSKNYHLENEVARLKKLVGERGSGSGSGSGSSLYPSLEDLKVDKVIQAQTAYSA<br>NPASQAFVLVDASAALPPDGNLYPKLYPELSQYMGLSLNEAEICESMPMVSGAPAQGQ<br>LVARPSSVNYMVAPVTGNDAGIRRAEIKHHHHHH*** |
| 12 | **mIL6ST-GCN4 LZ-Tfr Endosomal domain** | **MSAPRIWLAQALLFFLTTESIGQLLEPCGYIYPEFPVVQRGSNFTAICVLKEACLQHYYVN<br>ASYIVWKTNHAAVPREQVTVINRTTSSVTFTDVVLPSVQLTCNILSFGQIEQNVYGVTML<br>SGFPPDKPTNLTCIVNEGKNMLCQWDPGRETYLETNYTLKSEWATEKFPDCQSKHGTS<br>CMVSYMPTYYVNIEVWVEAENALGKVSSESINFDPVDKVKPTPPYNLSVTNSEELSSILK<br>LSWVSSGLGGLLDLKSDIQYRTKDASTWIQVPLEDTMSPRTSFTVQDLKPFTEYVFRIRSI<br>KDSGKGYWSDWSEEASGTTYEDRPSRPPSFWYKTNPSHGQEYRSVRLIWKALPLSEAN<br>GKILDYEVILTQSKSVSQTYTVTGTELTVNLTNDRYVASLAARNKVGKSAAAVLTIPSPHV<br>TAAYSVVNLKAFPKDNLLWVEWTPPPKPVSKYILEWCVLSENAPCVEDWQQEDATVN<br>RTHLRGRLLESKCYQITVTPVFATGPGGSESLKAYLKQAAPARGPTVRTKKVGKNEAVLA<br>WDQIPVDDQNGFIRNYSISYRTSVGKEMVVHVDSSHTEYTLSSLSSDTLYMVRMAAYT<br>DEGGKDGPEFTFTTPKFAQGEIEAIVVPVCLAFLLTTLLGVLFCFNKRDLIKKHIWPNVPD<br>PSKSHIAQWSPHTPPRHNFNSKDQRMKQLEDKVEELLSKNYHLENEVARLKKLVGERG<br>SGSGSGSGSMDQARSAFSNLFGGEPLSYTRFSLARQVDGDNSHVEMKLAVDEEENAD<br>NNTKANVTKPKHHHHHH*** |
| 13 | **hTNFR1** | **MGLSTVPDLLLPLVLLELLVGIYPSGVIGLVPHLGDREKRDSVCPQGKYIHPQNNSICCTK<br>CHKGTYLYNDCPGPGQDTDCRECESGSFTASENHLRHCLSCSKCRKEMGQVEISSCTVD<br>RDTVCGCRKNQYRHYWSENLFQCFNCSLCLNGTVHLSCQEKQNTVCTCHAGFFLRENE<br>CVSCSNCKKSLECTKLCLPQIENVKGTEDSGTTVLLPLVIFFGLCLLSLLFIGLMYRYQRWKS<br>KLYSIVCGKSTPEKEGELEGTTTKPLAPNPSFSPTPGFTPTLGFSPVPSSTFTSSSTYTPGDC<br>PNFAAPRREVAPPYQGADPILATALASDPIPNGGGGSGRVILEGGHHHHHH*** |
| 14 | **hTNFR1-Linker-Syndecan 1** | **MGLSTVPDLLLPLVLLELLVGIYPSGVIGLVPHLGDREKRDSVCPQGKYIHPQNNSICCTK<br>CHKGTYLYNDCPGPGQDTDCRECESGSFTASENHLRHCLSCSKCRKEMGQVEISSCTVD<br>RDTVCGCRKNQYRHYWSENLFQCFNCSLCLNGTVHLSCQEKQNTVCTCHAGFFLRENE<br>CVSCSNCKKSLECTKLCLPQIENVKGTEDSGTTVLLPLVIFFGLCLLSLLFIGLMYRYQRWKS<br>KLYSIVCGKSTPEKEGELEGTTTKPLAPNPSFSPTPGFTPTLGFSPVPSSTFTSSSTYTPGDC<br>PNFAAPRREVAPPYQGADPILATALASDPIPNGGGGSGRKDEGSYSLEEPKQANGGAY<br>QKPTKQEEFYALEGGHHHHHH*** |

| | | |
|---|---|---|
| 15 | **hTNFR1 C-term-CD63** | **MGLSTVPDLLLPLVLLELLVGIYPSGVIGLVPHLGDREKRDSVCPQGKYIHPQNNSICCTKCHKGTYLYNDCPGPGQDTDCRECESGSFTASENHLRHCLSCSKCRKEMGQVEISSCTVDRDTVCGCRKNQYRHYWSENLFQCFNCSLCLNGTVHLSCQEKQNTVCTCHAGFFLRENECVSCSNCKKSLECTKLCLPQIENVKGTEDSGTTVLLPLVIFFGLCLLSLLFIGLMYRYQRWKSKLYSIVCGKSTPEKEGELEGTTTKPLAPNPSFSPTPGFTPTLGFSPVPSSTFTSSSTYTPGDCPNFAAPRREVAPPYQGADPILATALASDPIPNGGGGSGRCCLVKSIRSGYEVMCCLLEGGHHHHHH*** |
| 16 | **hTNFR1 C-term-CD63 X 2** | **MGLSTVPDLLLPLVLLELLVGIYPSGVIGLVPHLGDREKRDSVCPQGKYIHPQNNSICCTKCHKGTYLYNDCPGPGQDTDCRECESGSFTASENHLRHCLSCSKCRKEMGQVEISSCTVDRDTVCGCRKNQYRHYWSENLFQCFNCSLCLNGTVHLSCQEKQNTVCTCHAGFFLRENECVSCSNCKKSLECTKLCLPQIENVKGTEDSGTTVLLPLVIFFGLCLLSLLFIGLMYRYQRWKSKLYSIVCGKSTPEKEGELEGTTTKPLAPNPSFSPTPGFTPTLGFSPVPSSTFTSSSTYTPGDCPNFAAPRREVAPPYQGADPILATALASDPIPNGGGGSGRCCLVKSIRSGYEVMCCLVKSIRSGYEVMLEGGHHHHHH*** |
| 17 | **hTNFR1 N-term-Syntenin** | **MGLSTVPDLLLPLVLLELLVGIYPSGVIGLVPHLGDREKRDSVCPQGKYIHPQNNSICCTKCHKGTYLYNDCPGPGQDTDCRECESGSFTASENHLRHCLSCSKCRKEMGQVEISSCTVDRDTVCGCRKNQYRHYWSENLFQCFNCSLCLNGTVHLSCQEKQNTVCTCHAGFFLRENECVSCSNCKKSLECTKLCLPQIENVKGTEDSGTTVLLPLVIFFGLCLLSLLFIGLMYRYQRWKSKLYSIVCGKSTPEKEGELEGTTTKPLAPNPSFSPTPGFTPTLGFSPVPSSTFTSSSTYTPGDCPNFAAPRREVAPPYQGADPILATALASDPIPNGGGGSGRSLYPSLEDLKVDKVIQAQTAFSANPANPAILSEASAPIPHDGNLYPRLYPELSQYMGLSLEGGHHHHHH*** |
| 18 | **hTNFR1 CD63** | **MAVEGGMKCVKFLLYVLLLAFCACAVGLIAIGVAVQVVLKQAITHETTAGSLLPVVIIAVGAFLFLVAFVGCCGACKENYCLMITFAIFLSLIMLVEVAVAIAGYVGGSRIHPSGVTGLVPSLGDREKRDSLCPQGKYVHSKNNSICCTKCHKGTYLVSDCPSPGRDTVCRECEKGTFTASQNYLRQCLSCKTCRKEMSQVEISPCQADKDTVCGCKENQFQRYLSETHFQCVDCSPCFNGTVTIPCKETQNTVCNCHAGFFLRESECVPCSHCKKNEECMKLCLPPPLANVTNPQDSGTEFGGIHTQGCVETIAIWLRKNILLVAAAALGIAFVEVLGIIFSCCLVKSIRSGYEVMDPPDLDN*** |
| 19 | **hTNFR1 delta MMP Intracellular-Foldon N-term-Syntenin** | **MGLSTVPDLLLPLVLLELLVGIYPSGVIGLVPHLGDREKRDSVCPQGKYIHPQNNSICCTKCHKGTYLYNDCPGPGQDTDCRECESGSFTASENHLRHCLSCSKCRKEMGQVEISSCTVDRDTVCGCRKNQYRHYWSENLFQCFNCSLCLNGTVHLSCQEKQNTVCTCHAGFFLRENECVSCSNCKKSLECTKLCLPQGTEDSGTTVLLPLVIFFGLCLLSLLFIGLMYRYQRWKGTGYIPEAPRDGQAYVRKDGEWVFLSTFLSPANGGGGSGRSLYPSLEDLKVDKVIQAQTAFSANPANPAILSEASAPIPHDGNLYPRLYPELSQYMGLSLEGGHHHHHH*** |
| 20 | **hTNFR1 delta MMP N-term-Syntenin** | **MGLSTVPDLLLPLVLLELLVGIYPSGVIGLVPHLGDREKRDSVCPQGKYIHPQNNSICCTKCHKGTYLYNDCPGPGQDTDCRECESGSFTASENHLRHCLSCSKCRKEMGQVEISSCTVDRDTVCGCRKNQYRHYWSENLFQCFNCSLCLNGTVHLSCQEKQNTVCTCHAGFFLRENECVSCSNCKKSLECTKLCLPQGTEDSGTTVLLPLVIFFGLCLLSLLFIGLMYRYQRWKNGGGGSGRSLYPSLEDLKVDKVIQAQTAFSANPANPAILSEASAPIPHDGNLYPRLYPELSQYMGLSLEGGHHHHHH*** |

| | | |
|---|---|---|
| 21 | **hTNFR1 delta MMP extracellular-Foldon N-term-Syntenin** | **MGLSTVPDLLLPLVLLELLVGIYPSGVIGLVPHLGDREKRDSVCPQGKYIHPQNNSICCTKCHKGTYLYNDCPGPGQDTDCRECESGSFTASENHLRHCLSCSKCRKEMGQVEISSCTVDRDTVCGCRKNQYRHYWSENLFQCFNCSLCLNGTVHLSCQEKQNTVCTCHAGFFLRENECVSCSNCKKSLECTKLCLNGGGGSGRGYIPEAPRDGQAYVRKDGEWVFLSTFLSPANGGGGSGRPQGTEDSGTTVLLPLVIFFGLCLLSLLFIGLMYRYQRWKGTNGGGGSGRGYIPEAPRDGQAYVRKDGEWVFLSTFLSPANGGGGSGRSLYPSLEDLKVDKVIQAQTAFSANPANPAILSEASAPIPHDGNLYPRLYPELSQYMGLSLEGGHHHHHH*MGLSTVPDLLLPLVLLELLVGIYPSGVIGLVPHLGDREKRDSVCPQGKYIHPQNNSICCTKCHKGTYLYNDCPGPGQDTDCRECESGSFTASENHLRHCLSCSKCRKEMGQVEISSCTVDRDTVCGCRKNQYRHYWSENLFQCFNCSLCLNGTVHLSCQEKQNTVCTCHAGFFLRENECVSCSNCKKSLECTKLCLNGGGGSGRGYIPEAPRDGQAYVRKDGEWVFLSTFLSPAPQGTEDSGTTVLLPLVIFFGLCLLSLLFIGLMYRYQRWKGTNGGGGSGRSLYPSLEDLKVDKVIQAQTAFSANPANPAILSEASAPIPHDGNLYPRLYPELSQYMGLSLEGGHHHHHHHHHHHH*** |
| 22 | **hTNFR1 delta MMP Extra and Intracellular-Foldon N-term-Syntenin** | **MGLSTVPDLLLPLVLLELLVGIYPSGVIGLVPHLGDREKRDSVCPQGKYIHPQNNSICCTKCHKGTYLYNDCPGPGQDTDCRECESGSFTASENHLRHCLSCSKCRKEMGQVEISSCTVDRDTVCGCRKNQYRHYWSENLFQCFNCSLCLNGTVHLSCQEKQNTVCTCHAGFFLRENECVSCSNCKKSLECTKLCLNGGGGSGRGYIPEAPRDGQAYVRKDGEWVFLSTFLSPANGGGGSGRPQGTEDSGTTVLLPLVIFFGLCLLSLLFIGLMYRYQRWKGTNGGGGSGRGYIPEAPRDGQAYVRKDGEWVFLSTFLSPANGGGGSGRSLYPSLEDLKVDKVIQAQTAFSANPANPAILSEASAPIPHDGNLYPRLYPELSQYMGLSLEGGHHHHHH*** |
